# Mesoamerica is a cradle and the Atlantic Forest is a museum of Neotropical butterfly diversity: Insights from the evolution and biogeography of Brassolini (Lepidoptera: Nymphalidae)

**DOI:** 10.1101/762393

**Authors:** Pável Matos-Maraví, Niklas Wahlberg, André V. L. Freitas, Phil DeVries, Alexandre Antonelli, Carla M. Penz

## Abstract

Regional species diversity is ultimately explained by speciation, extinction, and dispersal. Here we estimate dispersal and speciation rates of Neotropical butterflies to propose an explanation for their distribution and diversity of extant species. We focus on the tribe Brassolini (owl butterflies and allies): a Neotropical group that comprises 17 genera and 108 species, most of them endemic to rainforest biomes. We infer a robust species tree using the multispecies coalescent framework and a dataset including molecular and morphological characters. This formed the basis for three changes in Brassolini classification: 1) Naropina, SYN. NOV. is subsumed within Brassolina; 2) *Aponarope*, SYN. NOV. is subsumed within *Narope*; 3) *Selenophanes orgetorix*, COMB. NOV. is reassigned from *Catoblepia* to *Selenophanes*. By applying biogeographical stochastic mapping, we found contrasting species diversification and dispersal dynamics across rainforest biomes, which might be partly explained by the geological and environmental history of each bioregion. Our results reveal a mosaic of biome-specific evolutionary histories within the Neotropics, where butterfly species have diversified rapidly (cradles: Mesoamerica), have accumulated gradually (museums: Atlantic Forest), or have alternately diversified and accumulated (Amazonia). Our study contributes evidence from a major butterfly lineage that the Neotropics are a museum and cradle of species diversity.

## INTRODUCTION

Terrestrial biomes are heterogeneous in terms of species composition and distribution, and some areas have accumulated greater diversity and endemicity than others. A combination of three evolutionary mechanisms explains this biogeographical pattern: species origination, extinction, and dispersal (Goldberg *et al*., 2005; Jablonski *et al*., 2006). For example, a species-rich region can attain its present diversity by recent and exceptionally high rates of speciation, becoming a “cradle of diversity”, or by the persistence of old lineages via low species extinction rates, becoming a “museum of diversity” (e.g., Stebbins, 1974; Gaston & Blackburn, 1996; Bermingham & Dick, 2001; Richardson *et al*., 2001). Furthermore, a species-rich region might be a “source of diversity” when dispersal out of the region is high, or a “sink of diversity” when regional species diversity is accrued from external sources via dispersal into the region (Antonelli *et al*., 2018a). Characterizing the interplay among speciation, extinction, and dispersal is thus highly relevant for understanding the origin and evolution of total regional diversity and endemism (Wiens & Donoghue, 2004; Roy & Goldberg, 2007).

Much of the world’s biodiversity and endemism are concentrated in Neotropical rainforests in Central America, the Amazon basin, and the Atlantic Forest in southeastern Brazil. However, there is no consensus on whether these currently separated rainforests were connected once or multiple times, nor for how long (Jaramillo & Cárdenas, 2013). It is also unclear whether Neotropical rainforests represent centers of ecological stability that share a common evolutionary history (Moritz *et al*., 2000). Mammals and birds in Neotropical biodiversity hotspots, including the Brazilian Atlantic Forest and rainforests in Mesoamerica and the neighboring Chocó in the northwestern side of the Andes (Myers *et al*., 2000), apparently underwent rapid diversification (Igea & Tanentzap, 2019). In other groups, a larger number of lineages seem to have dispersed out of those areas compared to the rest of the Neotropics (e.g., Machado *et al*., 2018; Toussaint *et al*., 2019), suggesting that such rainforests are both cradles and sources of diversity (Igea & Tanentzap, 2019). Other Neotropical regions are, in contrast, museums of diversity given the old and gradual accumulation of species through time, such as Amazonia for Euptychiina (Nymphalidae) (Peña *et al*., 2010) and Troidini butterflies (Papilionidae) (Condamine *et al*., 2012). It thus seems that the checkered spatial assemblage of rainforest areas where speciation rates are high plus those where extinction rates are low might explain why the Neotropical region as a whole is considered both a cradle and museum of diversity (McKenna & Farrell, 2006; Moreau & Bell, 2013; Antonelli *et al*., 2018b). However, the impacts of geological and paleoenvironmental changes in the Neotropics together with the roles of speciation and dispersal have seldom been jointly elucidated among taxa occurring in individual rainforest areas.

Here we examine biogeographical patterns for butterflies in the tribe Brassolini (Nymphalidae: Satyrinae), an exclusively Neotropical monophyletic group (Freitas & Brown, 2004; Wahlberg *et al*., 2009; Espeland *et al*., 2018; Chazot *et al*., 2019a) consisting of 17 genera and 108 species found mainly in the rainforests of Mesoamerica, the northwestern slope (NW) of the Andes (including e.g., Chocó), the Atlantic Forest, and Amazonia (see Penz, 2007, for an overview). Studies that focused on individual brassoline genera used adult morphology (Penz, 2008, 2009a,b; Garzón-Orduña & Penz, 2009) and DNA data (Penz, Mohammadi, & Wahlberg, 2011a; Shirai *et al*., 2017) to infer species-level phylogenies, but several deep nodes had weak or even conflicting phylogenetic support in tribal-level analyses (Penz *et al*., 2013). We infer a time-calibrated species phylogeny of Brassolini by merging all previous datasets and generating new DNA and morphological data to infer the diversification and biogeographical histories of these butterflies. From a biogeographical perspective, if physical connection among Neotropical biomes existed during the Neogene and Quaternary periods (i.e., the past 23 Myr) (Hoorn *et al*., 2010), we expect that this will be reflected in the timing and magnitude of dispersal and speciation trends through time. To assess this expectation, we estimate dispersal rates and speciation for Brassolini butterflies.

## MATERIAL AND METHODS

### MOLECULAR DATASET

Adult specimens were collected in Mexico, Costa Rica, Ecuador, Peru, and Brazil (Supporting Information, Table S1), and identified to species based on various sources (Casagrande, 2002, 2004; Austin *et al*., 2007; Penz, 2008, 2009b,a; Garzón-Orduña & Penz, 2009; Penz *et al*., 2017, 2011a; Penz, Simonsen, & DeVries, 2011b; Chacón *et al*., 2012). Total DNA was obtained from two butterfly legs using QIAGEN’s DNeasy kits. Based on PCR primers and protocols described in Wahlberg & Wheat (2008), we Sanger-sequenced fragments of the mitochondrial COI gene (1,475 bp) and the nuclear genes CAD (850 bp), EF1α (1,240 bp), GAPDH (691 bp), RpS5 (617 bp), and *wingless* (400 bp) (outsourced to Macrogen, Seoul). Our molecular dataset included 64 species in 15 out of 17 Brassolini genera (Penz, 2007); the two missing genera (*Aponarope* and *Mielkella*) are monotypic. Specifically, we obtained genetic data from 83 individuals classified in 57 Brassolini species, and retrieved COI sequences available from the BOLD database (http://boldsystems.org) for an additional seven. The analyses also included genetic data of 19 outgroup taxa in the subfamily Satyrinae from public data deposited in GenBank. All DNA sequences from this study are in GenBank (accession numbers MK551348–MK551551).

**Table 1:** Secondary calibration points for divergence time estimation of Brassolini using BEAST v.2.6.3. We constrained the monophyly of representative taxa within the butterfly subfamily Satyrinae and used uniform age distributions based on Chazot *et al*. (2019a).

| MRCA (monophyletic) | Sampled taxa | Lower | Upper |
| --- | --- | --- | --- |
| Brassolini (Ingroup) - Morphini | Morphini: *Antirrhea*, *Morpho*, *Caerois* | 34 | 50 |
| Lethina - Parargina - Mycalesina | *Lethe*, *Pararge*, *Bicyclus* | 31 | 46 |
| Pronophilina - Euptychiina - Satyrina - Erebiina | *Calisto*, *Manerebia*, *Taygetis*, *Satyrus*, *Erebia* | 31 | 44 |
| Satyrini | Lethina, Parargina, Mycalesina, Pronophilina, Euptychiina, Satyrina, Erebiina, *Orsotriaena*, *Oressinoma* | 38 | 55 |
| Melanitini - Dirini | *Melanitis*, *Aeropetes* | 27 | 41 |
| Satyrinae (Root) | Brassolini, Morphini, Satyrini, Melanitini, Dirini, *Cithaerias*, *Pierella*, *Zeuxidia*, *Elymnias* | 46 | 65 |

### MORPHOLOGICAL AND TOTAL-EVIDENCE DATASET

Our morphological dataset included 255 discrete characters (201 from adults and 54 from early stages) for 64 species, 44 of them corresponding to the same species included in the molecular dataset. The morphological matrix included all 17 Brassolini genera and was concatenated from previous studies (Penz, 2007, 2008, 2009b,a; Garzón-Orduña & Penz, 2009; Penz *et al*., 2013) by eliminating duplicate characters and scoring new data for species missing entries. Specimens used were the same as listed in the original publications. The combined molecular and morphological dataset comprised all genera and 84 out of 108 Brassolini species (missing data coded as “?”). We call this the total-evidence dataset, which we used to infer a time-calibrated phylogeny.

### DISTRIBUTION DATASET AND BIOREGION DELIMITATION

We advocate for a data-informed bioregionalization of Brassolini geographical distribution. Specifically, we expect that parametric bioregionalization will better inform on the best-fit delimitation of areas for Brassolini compared to any arbitrary assignment of areas within the hierarchical classification by Morrone (2014a,b) (subregions, dominions or provinces).

We delineated bioregions (*sensu* Vilhena & Antonelli, 2015) using geo-referenced occurrence points of Brassolini identified with taxonomic certainty at the species level, of which 6,378 points came from GBIF (https://gbif.org; retrieved on 21^st^ January 2019) and 881 points from the ATLANTIC BUTTERFLIES dataset (Santos *et al*., 2018). Both databases follow the current taxonomic understanding of Brassolini, thus, we did not encounter dubious species names. We used the R v.3.5.3 (R Core Team, 2020) package CoordinateCleaner v.2.0-11 (Zizka *et al*., 2019) for automated cleaning of species occurrences. In addition to four *Caligo eurilochus* observations from tropical butterfly houses in Germany that were removed a priori, the software flagged 81 potential errors, such as occurrence records in the ocean, geographic coordinates identical to country centroids, and occurrences in close proximity to biodiversity institutions such as natural history museums (Supporting Information, Fig. S1). We uploaded the 7,174 vetted geo-referenced occurrences, all within the known natural geographical range of Brassolini (see below), to the web application Infomap Bioregions (Edler *et al*., 2017) at http://bioregions.mapequation.org/. We set the maximum and minimum values for cell sizes to 4 and 2, and for cell capacities to 40 and 4. We used different values of cluster cost, from 0.1 to 3.0, in steps of 0.1. The number of inferred areas remained stable between values of 1.8 and 2.4 and, thus, received the strongest support.

The bioregions identified correspond to formerly recognized biomes, or combinations of them: 1) Mesoamerica plus the NW flank of the Andes (including Chocó), 2) Brazilian Atlantic Forest, and 3) the remaining tropical South America, including the Amazon drainage basin and the seasonally dry diagonal biomes Cerrado and Caatinga (Fig. 1). Note that this delimitation based on different data and approach approximates the Dominions of Morrone (2014b): 1) a joint area including the Mesoamerican dominion plus the provinces of the Pacific dominion west of the Andes, in agreement with the strong floristic connection of Central America and Chocó (Pérez-Escobar *et al*., 2019), 2) the Parana dominion comprising the Brazilian Atlantic Forest, and 3) the remaining tropical provinces in South America (Fig. 1).

**Fig. 1:**
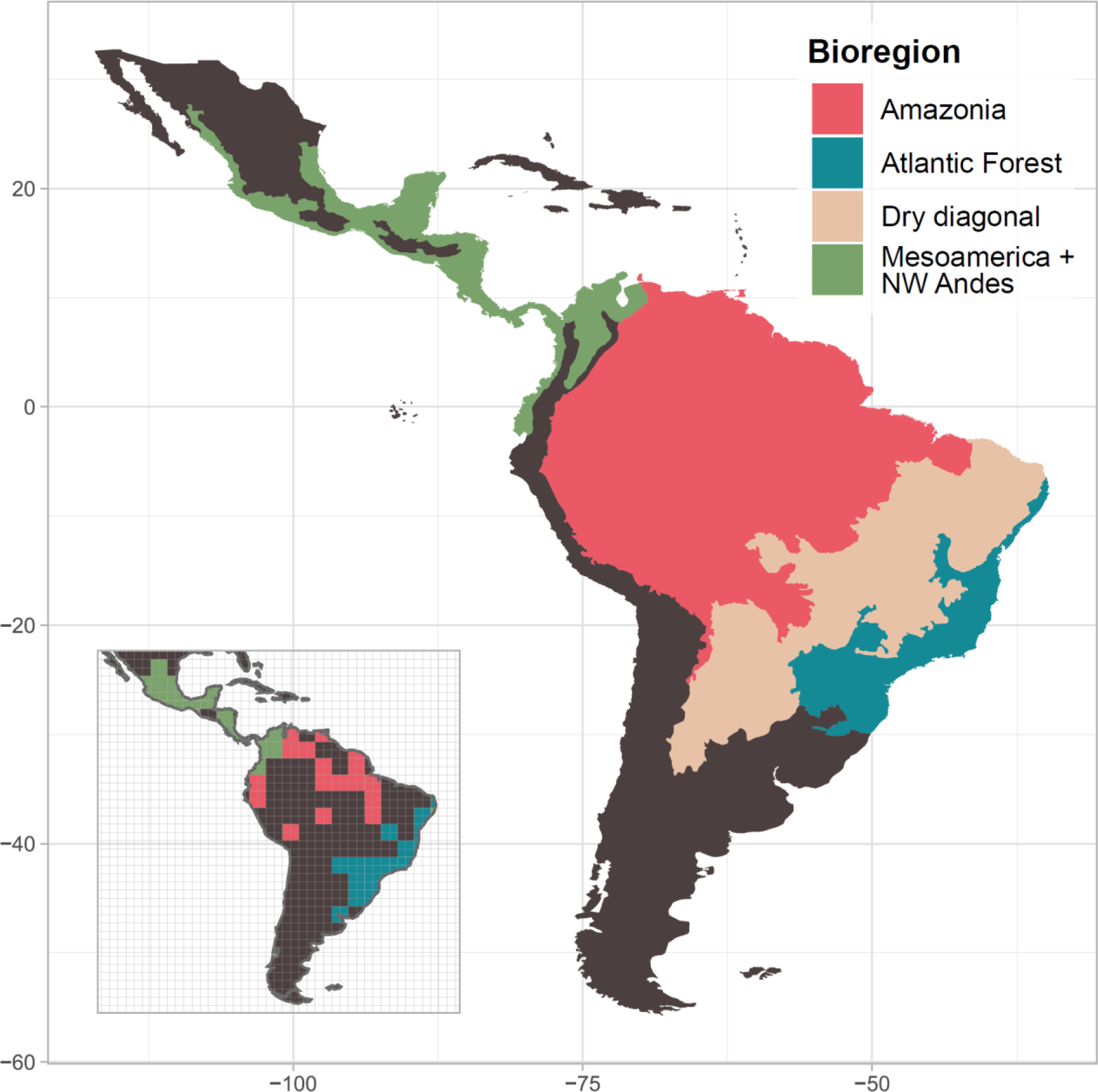
Map of the Neotropics and the defined bioregions based on occurrence data of Brassolini. Inset map shows the most stable bioregions under different cluster cost values by the software Infomap Bioregions using validated geo-coordinates from GBIF and ATLANTIC BUTTERFLIES databases. Note that the delineated bioregions approximate the Dominions of Morrone (2014b), encompassing the Mesoamerican plus Pacific dominions, except for three provinces east of the northern Andes; the Paraná dominion (Brazilian Atlantic Forest); the remaining dominions and provinces where Brassolini occurs. Shapefile from Löwenberg-Neto (2014).

To inform the biogeographical and within-area speciation rate analyses, we generated a species-level distributional dataset that includes all 108 valid Brassolini species. We surveyed the literature to score species occurrences in the defined bioregions because the geo-referenced dataset included only 72 Brassolini species with occurrence points. To investigate dispersal patterns across the South American dry diagonal, we kept this area separate (Fig. 1). We used distributional maps and observations, not always with primary geographic coordinates, from taxonomic revisions (Bristow, 1981, 1982, 1991; Casagrande, 2002; Furtado & Campos-Neto, 2004; Penz, 2008, 2009a,b; Garzón-Orduña & Penz, 2009; Penz *et al*., 2017) and published butterfly inventories in Mesoamerica (DeVries, 1983, 1994; Janzen & Hallwachs, 2009; Basset *et al*., 2015), Amazonia (Pereira Martins *et al*., 2017), Cerrado (Pinheiro & Emery, 2006; Emery *et al*., 2006; Silva *et al*., 2012; Pereira Martins *et al*., 2017; Dickens *et al*., 2019), Caatinga (Zacca & Bravo, 2012), and Atlantic Forest (Santos *et al*., 2011, 2018; Pérez *et al*., 2017; Melo *et al*., 2019; Soldati *et al*., 2019). We call this the presence/absence biogeographical dataset.

### PHYLOGENETIC INFERENCE AND DIVERGENCE TIME CALIBRATION

We ran preliminary phylogenetic analyses using each gene and morphological dataset to check for tree topology conflicts. The six molecular datasets were partitioned by codon position and the morphological dataset was partitioned by using homoplasy scores calculated through implied weighting parsimony (Rosa *et al*., 2019). We calculated homoplasy measurements *f* using TNT v.1.5 (Goloboff & Catalano, 2016) under the default concavity parameter *k* = 3, and we used these values to subdivide the morphological dataset into 11 partitions (Supporting Information, Table S3). All seven phylogenetic analyses (6 gene loci and morphology) were run using MrBayes v.3.2.6 (Ronquist *et al*., 2012) via the CIPRES Science Gateway v.3.3 (Miller *et al*., 2010). We performed model averaging over all substitution models within the GTR model family (Huelsenbeck *et al*., 2004), taking into account rate variation across sites (+I and +Γ models) for the molecular partitions, and we applied the Markov (MKv) substitution model (Lewis, 2001) for the morphological partitions. The Bayesian inferences were run two independent times for 50 million generations, sampling trees every 5,000^th^ generation and discarding the first 25% of the sampled distribution as burnin. After checking that the log-probabilities reached a stationary distribution, the average standard deviation of split frequencies were below 0.005, PSRF values close to 1.000, and the estimated sample sizes (ESS) above 200, we summarized the post-burnin sampled trees using the 50% majority-rule consensus method.

We inferred time-calibrated species trees using the StarBEAST2 v.0.15.5 package (Ogilvie *et al*., 2017) available in BEAST v.2.6.3 (Bouckaert *et al*., 2014). We removed the species *Opsiphanes camena* from the dataset because it had only 325 bp of the COI locus, and its phylogenetic position was not stable in the preliminary analyses. We used a gene-based partitioning strategy, each having unlinked tree models in the multispecies coalescent framework. We performed model averaging over 31 substitution models available in the bModelTest v.1.2.1 package (Bouckaert & Drummond, 2017). We applied uncorrelated relaxed-clock models for all partitions because strict molecular clock models were rejected based on the stepping-stone method (Table S4). We estimated two molecular clock rates, one for the mitochondrial locus and one for the nuclear loci, by using the gamma distribution for the clock rate priors (shape parameter (α) = 0.01 for mitochondrial clock and 0.001 for nuclear clock; scale parameter (β) = 1.0). We used the Yule tree model plus two molecular clocks because it received decisive support, based on path-sampling steps, over other models with different combinations of one, two, or six clocks, and the Yule and birth-death tree models (Table S5). We added the morphological dataset onto the species tree and we conditioned the characters to the Markov (Mk) substitution model (Lewis, 2001). BEAST2 assigns morphological partitions with unlinked site models based on characters having the same number of states. All morphological characters were considered to evolve under the same species tree and relaxed clock model with an uninformative rate prior (1/X distribution).

Inasmuch as there are no described fossils of Brassolini, we relied on secondary calibrations to time-calibrate the species tree. We conservatively used uniform distributions encompassing the 95% highest posterior density (HPD) from a large fossil-calibrated, genus-level butterfly phylogeny (Chazot *et al*., 2019a) to constrain six outgroup nodes (Table 1; Fig. S5).

The analyses were run four independent times for 300 million generations, via CIPRES. We sampled 20,000 trees from the posterior distribution and discarded the first 25% as burnin. We checked that the ESS values were above 200 using Tracer v.1.7.1 (Rambaut *et al*., 2018). We merged the independent runs using LogCombiner (part of the BEAST v.2.6.3 package) and we summarized the post-burnin species trees as a maximum clade credibility (MCC) tree in TreeAnnotator (part of the BEAST v.2.6.3 package).

As supplementary analyses, we inferred total-evidence phylogenetic trees using the concatenation approach assuming that all gene loci and morphological characters have evolved under the same tree topology. We carried out gene-tree discordance test to assess the convenience of the multispecies coalescent framework for our dataset. We performed likelihood-based tree topology tests to evaluate the agreement of the molecular data with three tree topologies: 1) the morphology-based systematics of Brassolini (Penz, 2007) suggesting that *Narope* is sister to remaining genera in the subtribe Brassolina, 2) the multispecies coalescent species tree, and 3) the concatenation-based tree topology (Supporting Information).

### WITHIN-AREA CLADOGENESIS AND DISPERSAL AMONG BIOREGIONS

We estimated ancestral geographic ranges and colonization rates among bioregions using the dispersal-extinction-cladogenesis (DEC) model (Ree *et al*., 2005) as implemented in the R package BioGeoBEARS v.1.1.2 (Matzke, 2013). We used the presence/absence biogeographical dataset and the four defined bioregions: 1) Mesoamerica plus NW Andes, 2) Atlantic Forest, 3) Amazonia, and the 4) Dry diagonal. The latter was kept separate to estimate dispersal between Atlantic Forest and Amazonia across a large area of less suitable environments for Brassolini. We did not constrain any *a priori* dispersal multiplier, nor did we stratify dispersal rates across the phylogeny. This allowed us to evaluate whether the data alone is in agreement with main geological and paleogeographical events in the Neotropics. To account for ancestral state uncertainty, we carried out 100 biogeographical stochastic mappings (BSM; Dupin *et al*., 2017). To account for divergence time uncertainty, we used 100 random topologies from the BEAST2 posterior distribution. To account for phylogenetic uncertainty of 25 missing species, we randomly added such lineages to their currently assigned genera in the 100 posterior phylogenies. In this case, we assumed that phylogenetic diversity per genus was maximized and we placed missing taxa in the crown genera given our comprehensive taxonomic sampling. All genera are monophyletic, but we conservatively joined *Selenophanes* and *Catoblepia* (posterior probability, PP = 1.0; Fig. 2) when assigning missing species because the monophyly of *Catoblepia* was only moderately supported (PP = 0.75). We also carried out the biogeographical analyses using the taxonomically incomplete inferred species trees to rule out any bias in the approach.

**Fig. 2:**
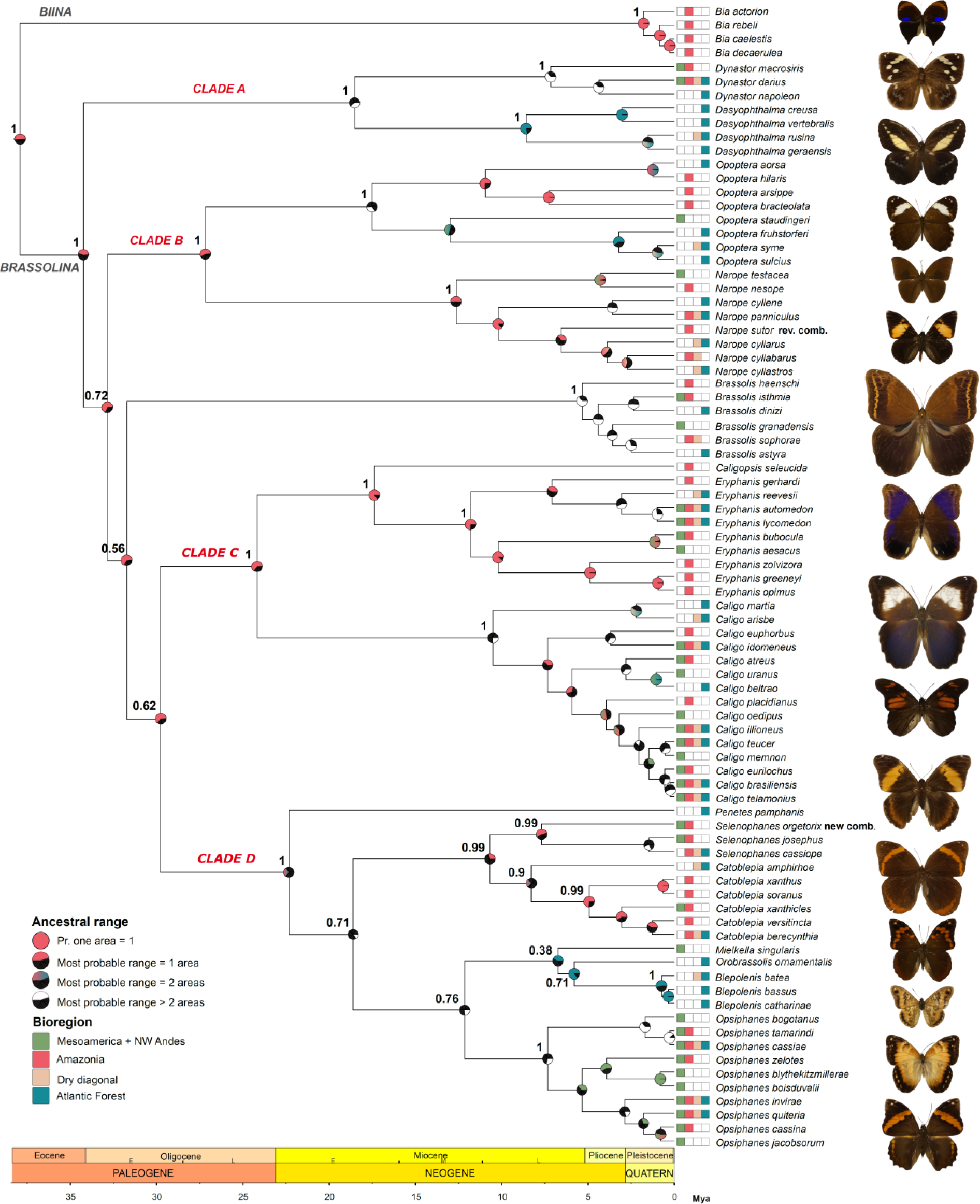
Time-calibrated species tree of the butterfly tribe Brassolini based on morphological and molecular data. The posterior probabilities for all genera and among genera are displayed next to the crown node. Ancestral range probability based on the DEC model and 10,000 biogeographical stochastic mappings is depicted as a pie chart following colors in legend. Images of representative Brassolini butterflies, all at the same scale: From top to down, *Bia actorion* (DeVries collection, PJD), *Dynastor darius* (Milwaukee Public Museum, MPM), *Dasyophthalma creusa* (MPM), *Opoptera fruhstorferi* (American Museum of the Natural History, AMNH), *Narope panniculus* (AMNH), *Brassolis astyra* (MPM), *Caligopsis seleucida* (PJD), *Eryphanis aesacus* (MPM), *Caligo martia* (MPM), *Penetes pamphanis* (MPM), *Selenophanes cassiope* (PJD), *Catoblepia berecynthia* (MPM), *Mielkella singularis* (AMNH), *Orobrassolis ornamentalis* (MPM), *Blepolenis bassus* (Museu de Zoologia - USP), *Opsiphanes quiteria* (PJD).

Variation in taxonomic opinion and effort across localities might be an important source of error in macroevolutionary analyses (Faurby *et al*., 2016). After analyzing trends in the year of description of each Brassolini species and the year of the latest revision of their current taxonomic status (e.g., when former subspecies were raised to species), we did not observe any taxonomic biases among Neotropical bioregions in terms of taxon discovery, description and taxonomic opinion (i.e., oversplitting species vs. multiple infraspecific taxa) (Supporting Information). We therefore estimated the number of dispersal events and within-area-cladogenesis events by simulating areas on nodes based on the 10,000 pseudoreplicated biogeographic histories (100 BSM × 100 posterior species trees). We calculated colonization rates through time as *c*_XtoY_(*t*_1_) = *d*_XtoY_(*t*_1_) / Br(*t*_1_), where *d*_XtoY_(*t*_1_) is the number of inferred dispersal events from area X to Y in a 2-million years interval, and Br(*t*_1_) is the total length of all branches in the 2-million-years interval (Antonelli *et al*., 2018a). In addition, we calculated the relative number of speciation events in area X through time as λ_X_(*t*_1_) = s_X_(*t*_1_) / L_X_(*t*_0_), where s_X_(*t*_1_) is the number of within-area cladogenesis in a 2-million-years interval, and L_X_(*t*_0_) is the number of inferred lineages in area X in the preceding 2-million-years interval (*t*_0_), calculated as the cumulative sum of within-area cladogenesis and dispersal into area X minus local extinction (Xing & Ree, 2017).

### SPECIATION WITHIN BIOREGIONS

For a second, independent estimate of within-area speciation, we calculated speciation rates for all tips and branches in the MCC species tree using BAMM v.2.5.0 (Rabosky, 2014). To account for the 25 missing species, we generated clade-specific sampling fractions at the genus level under the assumption of random taxonomic sampling (FitzJohn *et al*., 2009). We used the R package BAMMtools v.2.1.6 (Rabosky *et al*., 2014) to estimate prior values for the evolutionary rate parameters, and we assigned the expected number of shifts to 1.0, which is a conservative value that minimizes type I errors and is recommended for phylogenies including ∼100 species (Rabosky, 2014). We ran the analysis for 10 million generations, sampling event data every 1,000 generations. After discarding the first 20% samples, we checked that the ESS value was higher than 200 and the likelihood estimates reached a stationary distribution.

We used the function “dtRates” in BAMMtools to compute mean speciation rates along branches in ∼2-million years intervals (tau parameter set to 0.05, as the root of phylogeny was 37.9 Mya). We assigned the most probable inferred area from the 10,000 BSMs to the points where branches cross time intervals (Igea & Tanentzap, 2019). When the most probable area for a node and its parental one were the same, we assumed that the branch connecting both nodes occurred in such an area. When inferred areas for a node and its parental one were different, we assumed that the dispersal event occurred stochastically along the branch. We computed mean speciation rates only for cladogenesis events that most probably occurred in a single bioregion.

All the datasets and scripts used in this study can be found in TreeBASE (ID 25040) and Zenodo (DOI: http://doi.org/10.5281/zenodo.4500393).

## RESULTS

### PHYLOGENY AND DIVERGENCE TIMES

Our molecular dataset comprised 5,285 bp, of which 2,193 represented variable sites (Table 2). Our morphological dataset comprised 219 binary characters, 32 characters with 3 states, and 4 characters with 4 states. The multispecies coalescent model recovered four major clades with posterior probabilities (PP) of 1.0, namely, clade A, *Dynastor* and *Dasyophthalma*; clade B, *Opoptera* and *Narope*; clade C, *Caligopsis*, *Eryphanis*, and *Caligo*; clade D, *Penetes*, *Catoblepia*, *Selenophanes*, *Mielkella*, *Orobrassolis*, *Blepolenis*, and *Opsiphanes*. The maximum clade credibility (MCC) species tree depicts the following phylogenetic relationships: *Bia*: (clade A: (clade B: (*Brassolis*: (clade C: clade D)))), with divergence times among these clades between 29 and 34 Mya (entire highest posterior density HPD, 20.24–43.64 Mya). However, the relationships among clades B, C, D, and the genus *Brassolis* received low support (PP = 0.56– 0.72; Fig. 2). The concatenation-based analyses produced similar divergence times among major Brassolini clades compared to the multispecies coalescent species tree (between 26 and 30 Mya, entire HDP, 20.38–39.22 Mya) but recovered higher posterior probabilities for Brassolini deep nodes when using the molecular (0.97–1.00; Fig. S2) and total-evidence datasets (0.83–0.96; Fig. S3). Nevertheless, the consensus and MCC tree topologies among all concatenation-based phylogenetic analyses remained similar, except for the genus *Brassolis* as sister to the clade D, and clade C sister to these two. The likelihood-based tree topology tests could not resolve this main discrepancy between the multispecies coalescent and the concatenation-based species tree topologies (Table S6). The tree topology tests, however, clearly rejected the relationship of *Narope* as sister to the subtribe Brassolina suggested by previous cladistic analyses of morphological characters (*p*-value = 0.0003).

**Table 2:**
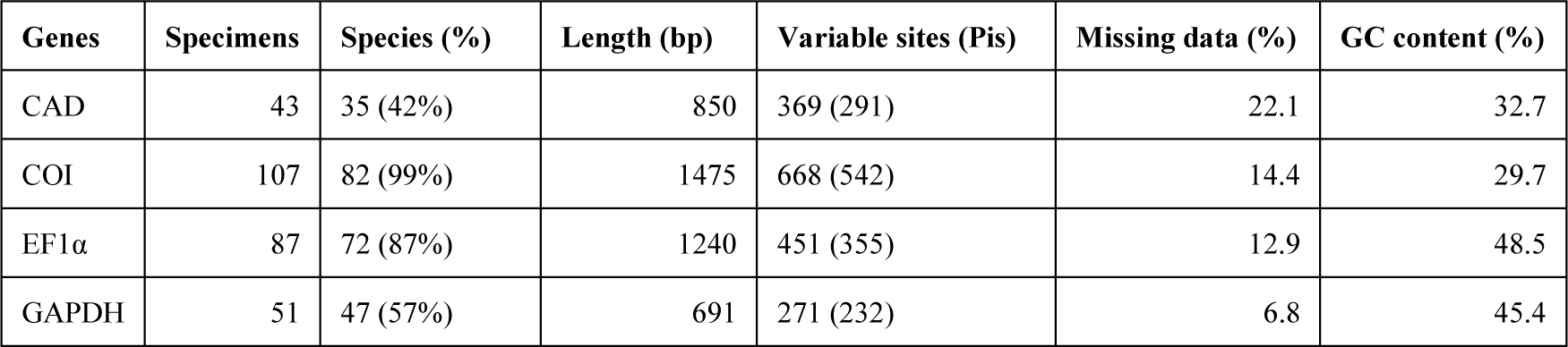
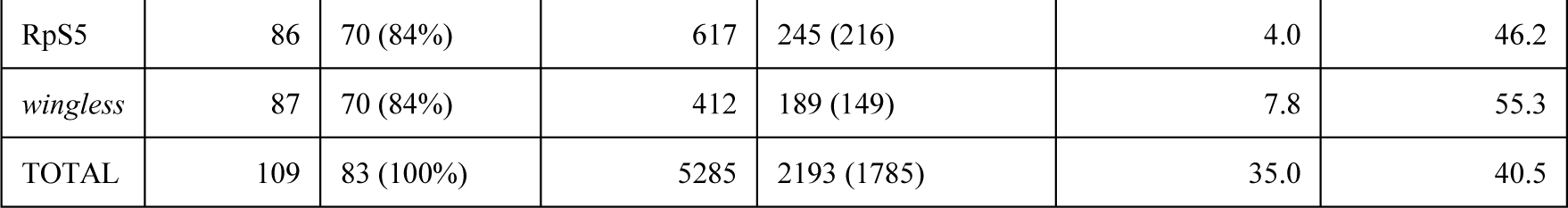
Characteristics of the multi-locus dataset. Specimens, species, and the molecular composition of the data matrix include outgroup taxa. (bp): base pairs; (Pis): number of parsimony-informative sites.

| Genes | Specimens | Species (%) | Length (bp) | Variable sites (Pis) | Missing data (%) | GC content (%) |
| --- | --- | --- | --- | --- | --- | --- |
| CAD | 43 | 35 (42%) | 850 | 369 (291) | 22.1 | 32.7 |
| COI | 107 | 82 (99%) | 1475 | 668 (542) | 14.4 | 29.7 |
| EF1α | 87 | 72 (87%) | 1240 | 451 (355) | 12.9 | 48.5 |
| GAPDH | 51 | 47 (57%) | 691 | 271 (232) | 6.8 | 45.4 |
| RpS5 | 86 | 70 (84%) | 617 | 245 (216) | 4.0 | 46.2 |
| *wingless* | 87 | 70 (84%) | 412 | 189 (149) | 7.8 | 55.3 |
| TOTAL | 109 | 83 (100%) | 5285 | 2193 (1785) | 35.0 | 40.5 |

Importantly, the total-evidence dataset supported posterior probability of 1.0 for the crown node of non-monotypic genera, regardless of phylogenetic method, except for two cases: *Narope* paraphyletic with respect to the monotypic *Aponarope*, and *Catoblepia* paraphyletic with respect to *Selenophanes* (see *Consequences for Brassolini systematics and classification* in the Discussion section).

### GEOGRAPHIC RANGE EVOLUTION AND COLONIZATION RATES

The inferred ranges coming from the 10,000 pseudoreplicated biogeographic histories and plotted against the MCC species tree in Fig. 2 suggested that Brassolini most probably (*Pr* = 0.50) originated in Amazonia at 38 Mya (entire HPD, 27.68–46.97 Mya). The early rapid radiation of Brassolini between 29 and 34 Mya also probably took place in Amazonia (*Pr* = 0.44–0.67).

When taking into account phylogenetic and divergence time uncertainties by using the Bayesian posterior distribution of Brassolini species trees, dispersal out of Amazonia had the highest rates compared to other bioregions (Fig. 3). Colonization rate from Amazonia to Mesoamerica showed a sharp increase by ∼2–4 Mya and is currently the highest rate among other dispersal types. Dispersal into Mesoamerica from other bioregions probably occurred throughout the Miocene but significantly increased at ∼6–8 Mya (Fig. S8). Dispersal out of Mesoamerica + NW Andes, however, was very low through time, and only since the past ∼2 Mya did it exponentially increase toward Amazonia. Dispersal from Amazonia to the dry diagonal increased throughout the Plio-Pleistocene and has since then equaled dispersal to Atlantic Forest. Dispersal from Amazonia into the Atlantic Forest was higher in the Oligocene–Miocene transition at ∼23–27 Mya than from/to any other Neotropical region. However, for most of the Miocene and Pliocene, dispersal into or out of the Atlantic Forest from/to any other bioregion was overall low through time, rendering the Atlantic Forest isolated; only since the Pleistocene ∼2 Mya dispersal into and from the dry diagonal has increased until the present. The dispersal patterns remain similar in supplementary biogeographical analyses using the taxonomically incomplete inferred species trees (Supporting Information).

**Fig. 3:**
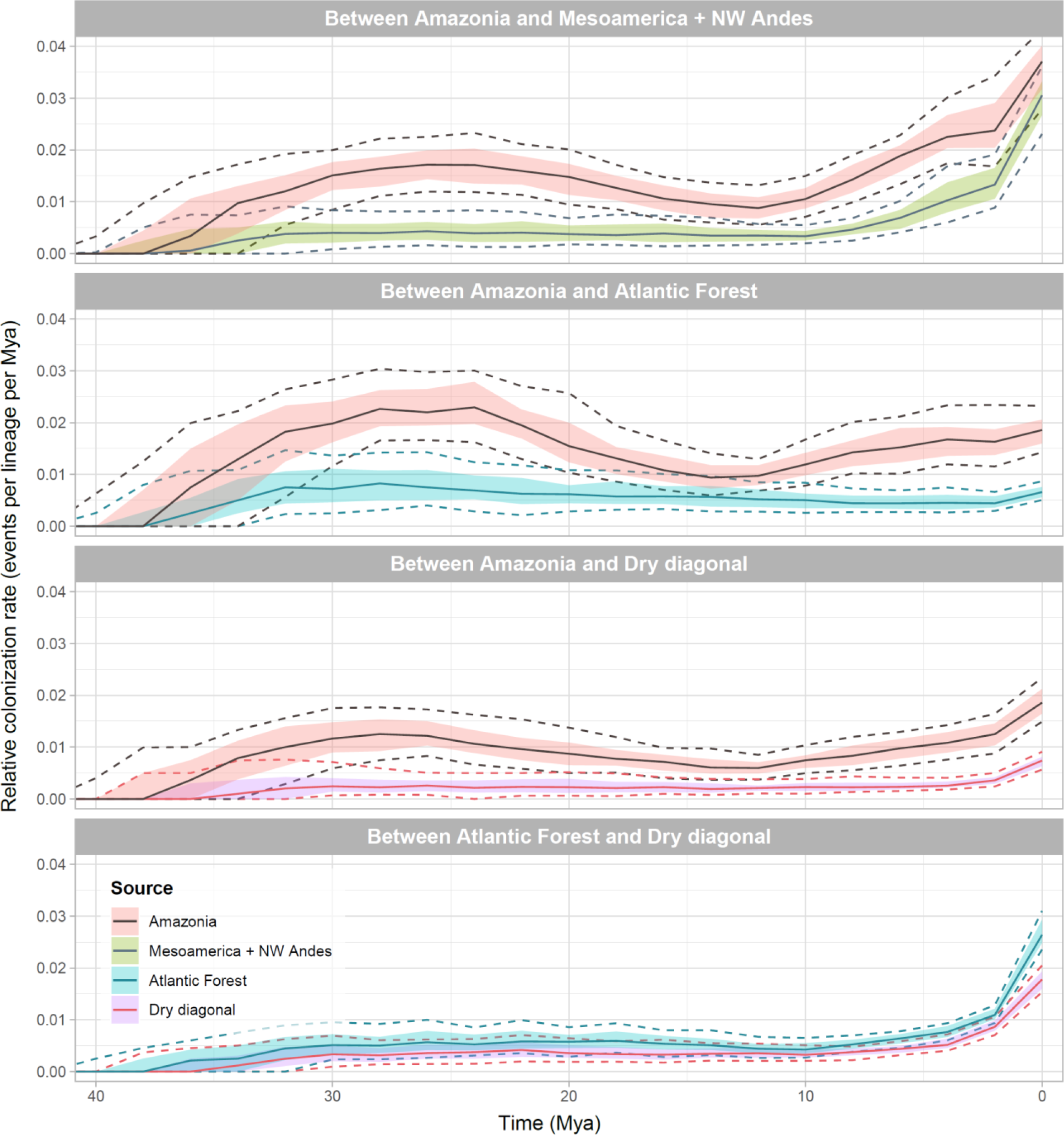
Colonization rates through-time based on 10,000 biogeographical stochastic mappings in BioGeoBEARS. Rates are displayed for a select pair of areas. Source areas (dispersal from) follow the color in the legend. Solid lines are the median values, colored ribbons are the lower and upper quartiles (0.25 and 0.75 quantiles), and dashed lines are the 0.1 and 0.9 quantiles.

### REGIONAL SPECIATION THROUGH TIME

The relative rates of within-area cladogenesis estimated in BioGeoBEARS and speciation estimated in BAMM were congruent (Fig. 4). Speciation rates increased through time in all areas except in the dry diagonal, which has had near zero endemic cladogenesis. Diversification in the Atlantic Forest might have begun during the Oligocene-Miocene transition at ∼20–24 Mya. During that time, the cladogenesis rate was slightly above 0.5 (Fig. 4A) and speciation rate began to increase (Fig. 4B), likely driven by the split of *Penetes* and, to a lesser extent, the stem lineage of *Dasyophthalma*. However, there seem to be a peak in cladogenesis at ∼8–10 Mya, specifically in the lineages comprising extant *Dasyophthalma*, *Orobrassolis* and *Blepolenis*. Afterwards, diversification in the Atlantic Forest remained constant throughout most of the Miocene until ∼2–4 Mya, when it slightly decreased towards the present. In contrast, speciation in Mesoamerica + NW Andes began later, at ∼8–12 Mya, remained constant during most of the Miocene and Pliocene, and sharply increased only at ∼2 to 4 Ma driven by the origin of extant endemic lineages within *Caligo*, *Opsiphanes*, and *Eryphanis*. Speciation in Amazonia was episodically higher than in the Atlantic Forest and Mesoamerica throughout most of the Miocene, but it also dramatically increased during the Pleistocene (∼2 Mya) by the split of several extant species within *Bia*, *Eryphanis*, and *Catoblepia*. At present, the speciation rate in Mesoamerica + NW Andes is similar to the Amazonian lineages, both apparently being higher than in the Atlantic Forest.

**Fig. 4:**
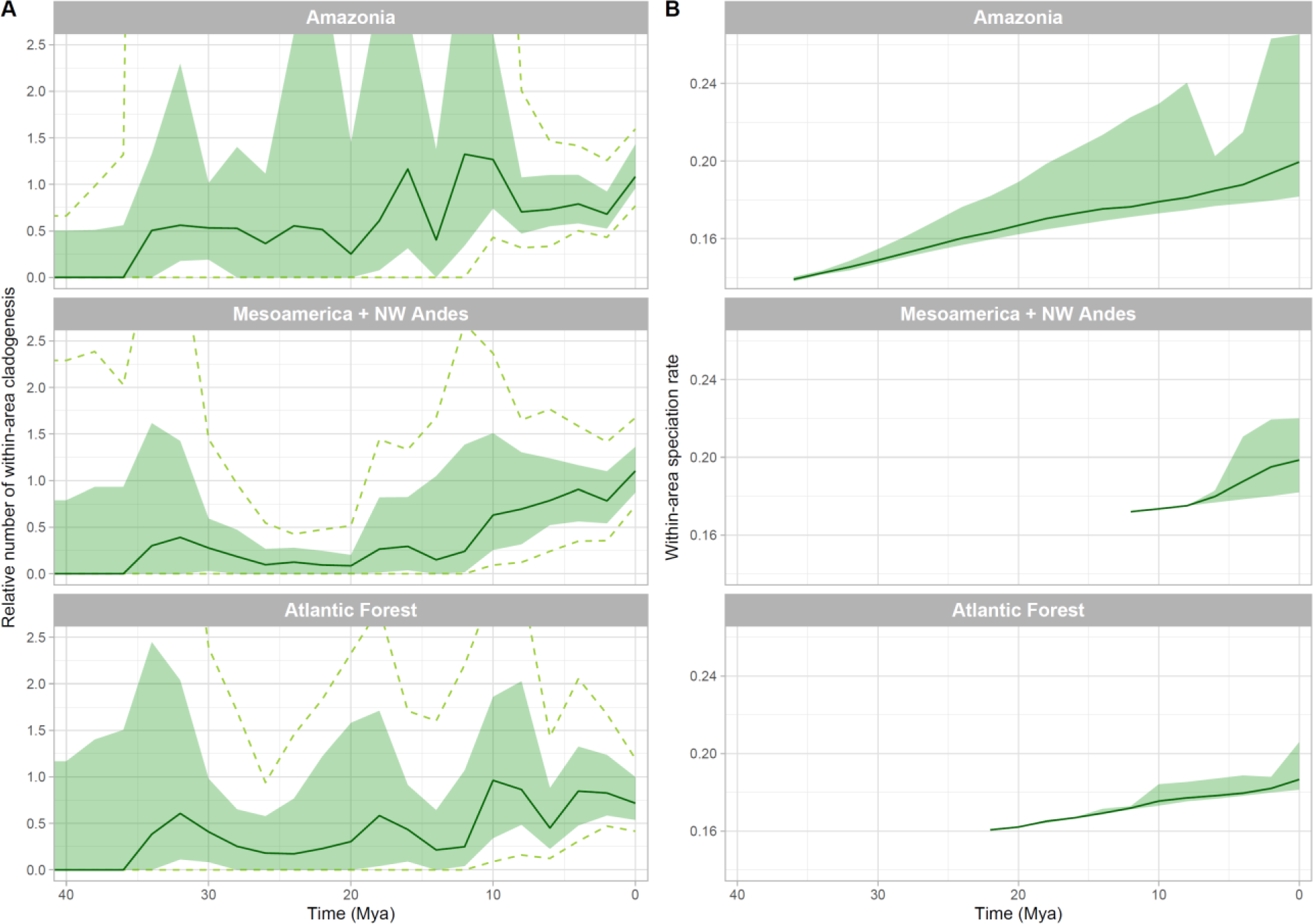
Within-area speciation through-time based on 100 posterior phylogenies and 100 biogeographical stochastic mappings (BSMs) in BioGeoBEARS. A) The relative number of within-area cladogenesis through time was calculated using a formula modified from Xing & Ree (2017) based on cladogenesis events occurring in one area. Solid lines represent median values, colored ribbons represent lower and upper quartiles (0.25 and 0.75 quantiles) and dashed lines represent 0.1 and 0.9 quantiles. B) Per-lineage speciation rates through time estimated by BAMM were assigned to a given bioregion based on the BSMs. Solid lines represent mean speciation rates per lineage and the colored ribbons the 5% and 95% confidence intervals.

## DISCUSSION

Here we investigated the evolutionary history of the butterfly tribe Brassolini, and studied the role of regional speciation and dispersal to understand the origin of extant species across Neotropical rainforest biomes (Fig. 5). By using molecular and morphological datasets and the multispecies coalescent framework, we propose a revised hypothesis of relationships among genera, and make three changes to the classification of Brassolini at the tribal and genus levels.

**Fig. 5:**
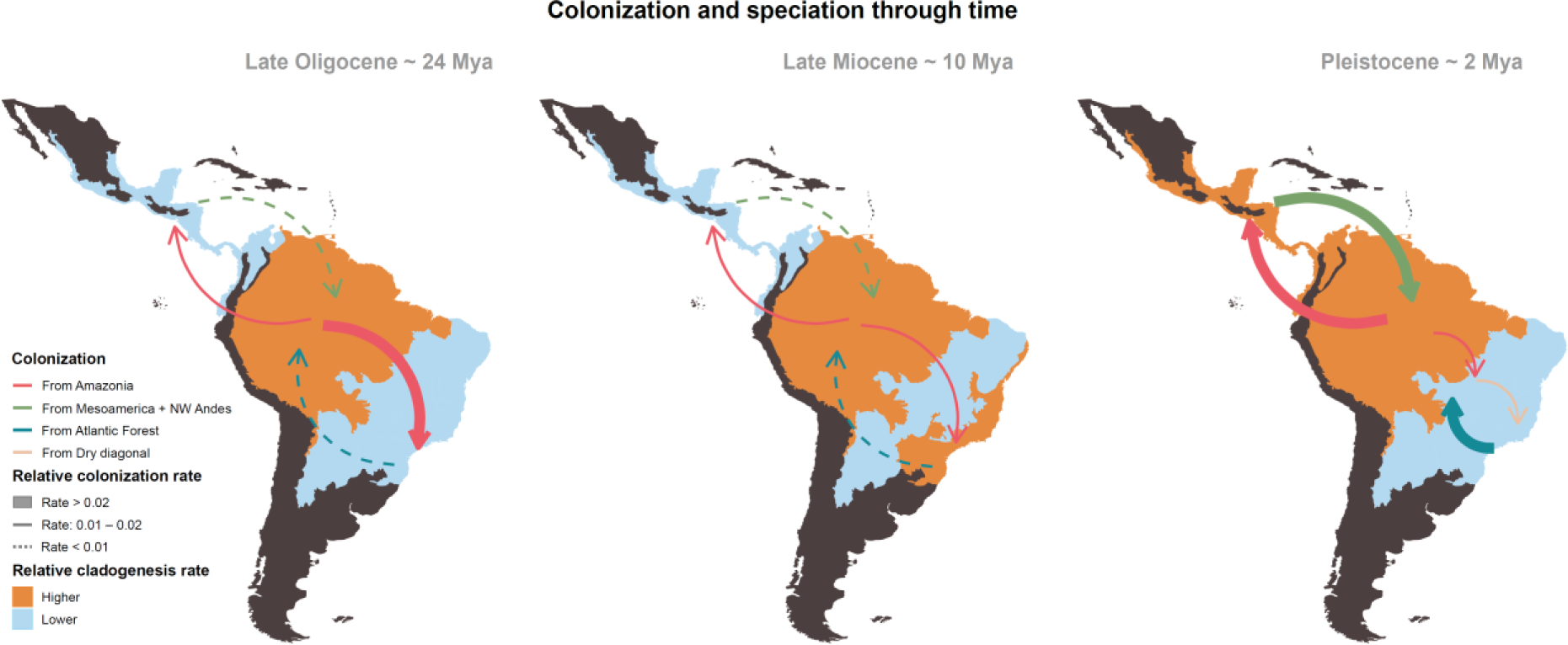
Summary of Brassolini colonization and speciation estimates in three time-windows, at 24, 10 and 2 Mya. Colored directional arrows represent dispersal from source areas, following colors in legend. The width and shape of arrows represent the estimated relative colonization rate between two areas in BioGeoBEARS, as depicted in Fig. 3. Bioregions were colored to qualitatively represent the estimated relative cladogenesis/speciation rates, as either center of speciation or area with comparatively lower speciation rates per time window.

### CONSEQUENCES FOR BRASSOLINI CLASSIFICATION

Brassolini has been previously subdivided in three subtribes based on shared morphological similarities: Biina (monotypic), Naropina (*Narope* and *Aponarope*), and Brassolina (remaining Brassolini genera) (Casagrande, 1995; Penz, 2007). Nonetheless, here *Narope* is sister to *Opoptera* (clade B in Fig. 2) with strong statistical support, and this clade is nested within the subtribe Brassolina (*sensu* Casagrande, 1995). These relationships have not been recovered in any cladistic analysis of Brassolini to date. At the genus level, the main discrepancy among the multispecies coalescent and the concatenation-based phylogenies was in the phylogenetic position of the genus *Brassolis*. Morphological character states of adult and early stages in this genus are highly polymorphic and divergent compared to other Brassolini genera, thus, obscuring the phylogenetic relationships within *Brassolis* and among this and other lineages (Garzón-Orduña & Penz, 2009; Penz *et al*., 2013). Nonetheless, the six molecular markers aided to resolve the apparent long-branch attraction between *Brassolis* and *Dynastor* seen in morphological phylogenies (Penz, 2007), and grouped *Brassolis* with clades C and D (*Caligo* and *Opsiphanes* groups) in both the multispecies coalescent and concatenation-based phylogenies.

We propose the following changes to the Brassolini classification:

1. The subtribe Naropina Stichel, 1925, **SYN. NOV.** is subsumed within Brassolina Boisduval, 1836, and Brassolini Boisduval, 1836, is redefined to include the subtribes Biina Herrich-Schäffer, 1864 (monotypic) and Brassolina.
2. The monotypic genus *Aponarope* Casagrande, 1982, **SYN. NOV.** is subsumed within *Narope* Doubleday, 1849. *Narope sutor* Stichel, 1916, was likely segregated in the monotypic *Aponarope* by Casagrande (1982) to accommodate the high number of autapomorphic characters. Using a cladistic analysis of 80 morphological characters, Penz *et al*. (2013) previously found that *Aponarope* is nested within *Narope* but conservatively retained *Aponarope* until more evidence increases the support for this relationship. Based on our results, we revise the genus assignment for *Narope sutor*, **COMB. REST.**, and synonymize *Aponarope*, **SYN. NOV.** to render the genus *Narope* monophyletic.
3. *Selenophanes orgetorix* (Hewitson, 1870), **COMB. NOV.** This species grouped with members of *Selenophanes* Staudinger, 1887, with high posterior probability (0.99) in our analysis, which justifies the revised generic assignment. Using COI barcoding sequences, Shirai *et al*. (2017) also found that the genus *Catoblepia* Stichel, 1902, was paraphyletic with respect to *Selenophanes* due to the sister relationship between *S. orgetorix* and *Selenophanes* species.

### SPECIES DIVERSIFICATION AND DISPERSAL

One third of Brassolini species are endemic to rainforests in Mesoamerica, NW slope of the Andes including the Chocó, and the Brazilian Atlantic Forest. Rapid butterfly speciation has probably occurred in Mesoamerica and NW Andes since the Pleistocene, suggesting that it acted as a cradle of diversity. In contrast, low and constant diversification since at least the early Miocene has shaped Brassolini endemic diversity in the Atlantic Forest. This finding, coupled with the survival of old lineages found mainly in montane habitats (e.g., *Penetes*, *Dasyophthalma*, *Orobrassolis*) suggests that the Atlantic Forest acts as a museum of diversity for Brassolini. Our results also suggest that Brassolini diversified mainly in Amazonia through episodic increases in cladogenesis rate throughout the Miocene from about 20 Ma (Fig. 4A), and such diversification events might have been accompanied by increasing dispersal out of this bioregion.

The diversification and biogeographical history of Brassolini are consistent with the source, museum, and cradle of diversity scenarios previously proposed for the Amazon rainforest. Amazonia has been the main source of Neotropical vertebrate and plant species diversity by means of dispersal during the past 60 million years (Antonelli *et al*., 2018a). Rapid species diversification of some butterfly groups occurred there since at least the Oligocene (e.g., Matos-Maraví *et al*., 2013; Chazot *et al*., 2016; Toussaint *et al*., 2019), and old lineages might have gradually accumulated in Amazonia since that time (Peña *et al*., 2010; Condamine *et al*., 2012). However, more recent diversification episodes supporting Amazonia as alternating museum and cradle of diversity could have been triggered by paleoenvironmental changes during the Miocene (Antonelli *et al*., 2009; Antonelli & Sanmartín, 2011; Chazot *et al*., 2019b).

The establishment of the Pebas wetland system in western Amazonia during most of the Miocene (∼8–24 Mya) (Wesselingh & Salo, 2006) has likely hindered dispersal from Amazonia toward the NW Andes and the Pacific in some butterfly groups (Elias *et al*., 2009; De-Silva *et al*., 2016, 2017; Chazot *et al*., 2019b). In the case of Brassolini, the Pebas wetland system has also likely hindered dispersal out of Amazonia (Fig. 3) but it might not have drastically limited diversification (Fig 4). This wetland system was occasionally affected by marine flooding events, which were probably short-lived, and at least two dated events at ∼13.7 and ∼17.8 Mya (Jaramillo *et al*., 2017a) were concurrent with the early and rapid cladogenesis events of major Brassolini clades in Amazonia. The Pebas wetland might in fact have increased opportunities for dispersal across Amazonia and diversification of some typical coastal plants (Bernal *et al*., 2019), including many of the Brassolini’s hostplants in the region: Arecaceae (palms), Poaceae (bamboos), and Zingiberales such as Marantaceae (Beccaloni *et al*., 2008; Janzen *et al*., 2009). It is plausible that the evolutionary consequences of the Miocene marine incursions in western Amazonia might have been mixed and rather ephemeral (Musher *et al*., 2019), restricting dispersal and diversification in some butterfly lineages such as Ithomiini (Chazot *et al*., 2019b) while providing ecological opportunities for diversification of others, such as Brassolini.

The increase in diversification of Amazonian lineages in the Pleistocene might be related to younger paleoenvironmental changes, such as fluctuating river dynamics mediated by Quaternary glacial cycles (Ribas *et al*., 2012). For example, biogeographical structure in the genus *Bia* seems to be associated with major Amazon-basin rivers (Penz *et al*., 2017), suggesting that riverine barriers might have been involved in allopatric differentiation in this genus. Indeed, in some areas surrounding the Amazon River, the distance between river banks is much larger than the estimated daily dispersal of species such as *Bia actorion* (Tufto *et al*., 2012; Penz *et al*., 2015). Other hypotheses for the diversification of Neotropical butterflies in the Pleistocene have been put forward, such as rainforests acting as refugia (Garzón-Orduña *et al*., 2014), but this has not been corroborated yet in a macroevolutionary framework (Matos-Maraví, 2016). A phylogeographic evaluation including more samples per species would be necessary to estimate the influence of river barriers or rainforest refugia in differentiating Brassolini populations, and potentially to explain the apparent increase in diversification of Amazonian butterflies during the Pleistocene.

Miocene expansions of wet forests between the Atlantic Forest and Amazonia across the dry diagonal might have been important determinants of ancient Brassolini diversity. The onset of a period of global cooling at ∼10 to 15 Mya likely drove the expansion of open environments (Simon *et al*., 2009; Azevedo *et al*., 2020), but intermittent forest connections between Amazonia and the Atlantic Forest have been documented based on palynological data and biogeographic studies (e.g., Costa, 2003; Werneck, 2011; Prates *et al*., 2016; Trujillo-Arias *et al*., 2017). Although evidence of repeated Neogene connection between the Atlantic Forest and Amazonia is still scarce and restricted to phylogeographic studies (e.g., Ledo & Colli, 2017; Capurucho *et al*., 2018), the origin of the modern transcontinental Amazon river by ∼10 Mya might have facilitated dispersal of rainforest taxa (Albert *et al*., 2018). Indeed, we found that late Miocene dispersal rates between these wet forest biomes slightly increased through time. In contrast, the Atlantic Forest has recently become a source of diversity for present dry diagonal as Brassolini dispersal sharply increased during the Pleistocene, in agreement with previous phylogeographic evidence (Costa, 2003; Batalha-Filho *et al*., 2013; Thomé *et al*., 2016). We suggest that episodic Miocene wet forest expansions might have represented ecological opportunities for Brassolini to disperse across Central Brazil, either by increased sizes of wet forest galleries in the dry diagonal biome, or by fully connecting Amazonia and Atlantic Forest.

By the Plio-Pleistocene, speciation in Mesoamerica and NW Andes seems to have increased and surpassed the speciation rate in the Atlantic Forest. Biotic dispersal into Mesoamerica from South America (e.g., Mullen *et al*., 2011; Bacon *et al*., 2015) and butterfly dispersal from Central America to other Neotropical areas (Toussaint *et al*., 2019) likely occurred throughout most of the Miocene facilitated by land availability, with Uranium-lead dating confirming that segments of the Panama arc had already emerged by 13–15 Mya (Montes *et al*., 2015). However, only at the end of the Pliocene ∼3 to 4 Mya did shallow seawaters fully recede from the region (Coates *et al*., 2004; Montes *et al*., 2012; O’Dea *et al*., 2016; Jaramillo *et al*., 2017b), thus increasing opportunities for terrestrial species diversification in lowland areas that were sporadically inundated by seawater during the Miocene (Hosner *et al*., 2016). In contrast, speciation in the Atlantic Forest remained low and dispersal out of this region decreased until the mid-Pleistocene. The contraction of the Atlantic Forest since the mid-Miocene (Simon *et al*., 2009; Edwards *et al*., 2010; Strömberg, 2011; Werneck, 2011) may have restricted Brassolini speciation to montane habitats in southeastern Brazil where most extant species occur, and old lineages having low speciation rates through time survived in remnants of rainforest.

### FUTURE DIRECTIONS

The evolutionary patterns inferred here indicate that geography has been an important determinant of the diversity and distribution of extant Brassolini butterflies. Other unmeasured or unknown traits might have driven the observed disparate trends in speciation through time, for example larval hostplants. However, many Mesoamerican Brassolini feed on the same plant families (e.g., Arecaceae), and even genera (e.g., *Euterpe*, *Bactris*), as do Atlantic Forest taxa (Beccaloni *et al*., 2008; Janzen *et al*., 2009; Robinson *et al*., 2010). Thus, despite the limited data at hand, it seems that larval hostplants might have contributed little to the disparate diversification dynamics of Brassolini.

We are aware that estimating diversification rates shifts through time is challenging (Moore *et al*., 2016). However, further evaluation via simulated data suggested that BAMM can infer speciation rates accurately from extant-taxon phylogenies (Rabosky *et al*., 2017). Although low extinction rates might explain the museum of diversity pattern, we have conservatively focused only on regional species origination rates to explain lineage accumulation within bioregions. Furthermore, (Louca & Pennell, 2020) stressed that the identifiability of diversification processes is problematic in a model-selection approach, but this remains to be evaluated in methods that use rate shift models such as BAMM (Laudanno *et al*., 2020). Reassuringly, we also used a fundamentally distinct measure of within-area cladogenesis (Xing & Ree, 2017) that corroborated the pattern of recent and rapid speciation in Mesoamerica + NW Andes, and an older lineage accumulation and lower diversification in the Atlantic Forest. However, our proposed checkered assemblage of Neotropical cradles and museums certainly needs to be evaluated with other larger phylogenies of rainforest lineages, particularly in light of natural history differences among various butterfly groups.

Our approach to disentangle regional speciation from dispersal to understand the origin of within-area species diversity has two strengths. First, we defined bioregions that fit the distribution and composition of Brassolini species by using geo-referenced occurrences, rather than assuming any subjective criteria to adopt areas from hierarchical bioregion classifications (Vilhena & Antonelli, 2015). This data-driven approach revealed a clustering of butterfly communities from southern North America to Mesoamerica and the NW side of the Andes, which conforms with the likely emergence of landmass in Central America, Chocó and northeastern Colombia after the collision of the Panama Block and northwestern South America by the late Oligocene (Coates & Stallard, 2013; Jaramillo, 2018). Furthermore, the inference of large bioregions for Brassolini agrees with its apparent poor spatial structuring and high dispersal capabilities. For instance, given a suitable environment, ancestral *Bia* had the capability to disperse across pan-Amazonia in only 1,463–3,115 years (Penz *et al*., 2015). The delimitation of Mesoamerica + NW Andes and the Atlantic Forest as separate bioregions has likely been determined by ecological and habitat suitability rather than geographical barriers to Brassolini dispersal; this being a similar biogeographical pattern found in other Neotropical rainforest taxa (Dexter *et al*., 2017; Pérez-Escobar *et al*., 2019).

## CONCLUSIONS

We investigated whether the diversification and biogeographical history of butterflies classified in the tribe Brassolini conforms to the premise that the Neotropics consists of a regional network of museums and cradles of diversity (Fig. 5). We found that endemic species to Mesoamerica plus the NW side of Andes have originated by old dispersal events into the region, and that speciation rates increased only at about 2 Mya (cradle of diversity). In contrast, endemic species to the Brazilian Atlantic Forest likely arose under low and continuous speciation rates since at least 12 Mya (museum of diversity), and was likely intermittently connected to Amazonia since the Miocene. The dynamic evolutionary history of Amazonian Brassolini lineages alternately diversifying and accumulating old lineages appears to reflect paleoenvironmental changes, including landscape reconfigurations in western Amazonia and paleoclimate fluctuations during the Neogene and Quaternary. By focusing on a representative lineage of a major insect order, our study paves the way for understanding why the Neotropics should be considered both a museum and cradle of diversity.

## ACKNOWLEDGEMENTS

We are grateful to Carlos Peña, Mirna Casagrande, Helena Romanowski, Augusto H. B. Rosa, and Andrew Warren for contributing specimens used in this study; and to Diana Silva, Gerardo Lamas, and SERFOR (Ministerio de Agricultura, Peru) for assistance with research permits (No Autorización 223-2017-SERFOR/DGGSPFFS). We thank the anonymous reviewers for their helpful comments. We are also grateful to the ICMBio for the research permits (SISBIO n° 10802-5, 10802-24 and 53016-10). Brazilian specimens are registered under the SISGEN (A16771F). We acknowledge the computational resources provided by the CESNET LM2015042 and the CERIT Scientific Cloud LM2015085, provided under the programme “Projects of Large Research, Development, and Innovations Infrastructures”. This publication is part of the RedeLep ‘Rede Nacional de Pesquisa e Conservação de Lepidópteros’ SISBIOTA-Brasil/CNPq (563332/2010-7). PMM was supported by a Marie Skłodowska-Curie fellowship (European Commission, Grant No. MARIPOSAS-704035), by the Grant Agency of the Czech Republic (GAČR grant: GJ20-18566Y), and by a fellowship from the PPLZ programme (Czech Academy of Sciences, Fellowship No. L200961951). AVLF acknowledges support from FAPESP (2011/50225-3, 2012/50260-6, 2013/50297-0), from the Brazilian Research Council - CNPq (303834/2015-3), from the National Science Foundation (DEB-1256742) and from the United States Agency for International Development - USAID / the U.S. National Academy of Sciences (NAS), under the PEER program (Sponsor Grant Award Number: AID-OAA-A-11-00012) (Mapping and Conserving Butterfly Biodiversity in the Brazilian Amazon). NW acknowledges support from the Swedish Research Council. AA acknowledges support from the Swedish Research Council, the Swedish Foundation for Strategic Research, the Knut and Alice Wallenberg Foundation, and the Royal Botanic Gardens, Kew.

## SUPPORTING INFORMATION

Additional Supporting Information may be found in the online version of this article at the publisher’s website:

**Supplementary data analyses** (pp. 2–12).

**Figure S1.** Map with the clean and flagged Brassolini geo-referenced occurrences (pp. 13).

**Figure S2.** Consensus trees of molecular and morphological datasets (pp. 14–22).

**Figure S3.** Consensus tree of concatenated total-evidence dataset (pp. 23–24).

**Figure S4.** Parsimony-based partitioned Bremer support scores (pp. 25–26).

**Figure S5.** Time-calibrated tree using the multispecies coalescent (pp. 27–28).

**Figure S6.** Brassolini taxonomic resolution across the Neotropics (pp. 29–30).

**Figure S7.** Ancestral range probabilities plotted on species tree (pp. 31–35).

**Figure S8.** Dispersal rate through time between bioregions (pp. 36–38).

**Figure S9.** Within-area cladogenesis events through time (pp. 39–41).

**Table S1.** Voucher locality information and associated genetic data (pp. 42, separate file).

**Table S2.** Best-fit partitioning scheme for the molecular dataset (pp. 42).

**Table S3.** Best-fit partitioning scheme for the morphological dataset (pp. 43–44).

**Table S4.** Bayes factor between the strict and relaxed clock models (pp. 45).

**Table S5.** Bayes factor among clock partitions and tree models (pp. 46).

**Table S6.** Tree topology test of early divergent Brassolini lineages (pp. 47).

**Table S7.** Sampling fractions for taking into account missing species (pp. 48).

