## Supporting Information for "Mesoamerica is a cradle and the Atlantic Forest is a museum of Neotropical butterfly diversity: Insights from the evolution and biogeography of Brassolini (Lepidoptera: Nymphalidae)"

### Table of Contents:


*Supplementary data analyses* ……………. 2

Figure S1: *Map with the clean and flagged Brassolini occurrences* ……………. 13

Figure S2: *Consensus trees of molecular and morphological datasets* ……………. 14

Figure S3: *Consensus tree of concatenated total-evidence dataset* ……………. 23

Figure S4: *Parsimony-based partitioned Bremer support scores* ……………. 25

Figure S5: *Time-calibrated tree using the multispecies coalescent* ……………. 27

Figure S6: *Brassolini taxonomic resolution across the Neotropics* ……………. 29

Figure S7: *Ancestral range probabilities plotted on species tree* ……………. 31

Figure S8: *Dispersal rate through time between bioregions* ……………. 36

Figure S9: *Within-area cladogenesis events through time* ……………. 39

Table S1: *Voucher locality information and associated genetic data* ……………. 42

Table S2: *Best-fit partitioning scheme for the molecular dataset* ……………. 42

Table S3: *Best-fit partitioning scheme for the morphological dataset* ……………. 43

Table S4: *Bayes factor between the strict and relaxed clock models* ……………. 45

Table S5: *Bayes factor among clock partitions and tree models* ……………. 46

Table S6: *Tree topology test of early divergent Brassolini lineages* ……………. 47

Table S7: *Sampling fractions for taking into account missing species* ……………. 48

### Phylogenetic inference and divergence time calibration

*Automated cleaning geographic coordinates*

We used the R v.3.5.3 (R Core Team, 2019) package CoordinateCleaner v.2.0-11 (Zizka *et al.*, 2019) to flag potential errors in the GBIF and ATLANTIC BUTTERFLIES Brassolini datasets. We carried out several tests including identifying records with geographic coordinates falling in the ocean, country capitals, country centroids, GBIF headquarters, biodiversity institutions such as natural history museums, and invalid/equal latitude and longitude coordinates. In total, 81 occurrences were flagged and excluded from our bioregion delimitation analyses. A map depicting the 7,174 cleaned and 81 flagged occurrences is shown in Fig. S1.

*Concatenated molecular data*

We ran a concatenated phylogenetic analysis using the Brassolini molecular dataset to evaluate any major tree topology difference from the multispecies coalescent species tree. We estimated the best-fit partition strategy for the multi-locus dataset using PartitionFinder v2.1.1 (Lanfear *et al.*, 2017). The program was run with 18 data blocks, one for each codon position in all six genes, and we used the greedy search option. The linked branch lengths option is preferred over the unlinked branches based on Bayesian Information Criterion value ($BIC_{linked}$ = 121,422.6; $BIC_{unlinked}$ = 124,848.4). The best partition scheme consists of 8 subsets (Table S2).

The phylogenetic analysis was conducted in MrBayes v.3.2.6 (Ronquist *et al.*, 2012) via the CIPRES Science Gateway v.3.3 (Miller *et al.*, 2010). We performed model averaging over all substitution models within the GTR family, using a reversible jump MCMC (Huelsenbeck *et al.*, 2004). The analysis took into account rate variation across sites by using the +I and +Γ models, and was run two independent times for 50 million generations. We sampled 5,000 trees from the posterior distribution and we discarded the first 25% of sampled trees as burnin. We checked the mixing of chains in both independent runs by inspecting that the log-probabilities reached a stationary distribution, the average standard deviation of split frequencies were below 0.005, PSRF values close to 1.000, and the estimated sample sizes (ESS) above 200. We summarized the post-burnin sampled trees using the 50% majority-rule consensus method (Fig. S2).

*Concatenated total-evidence data*

We removed the DNA sequence of *Opsiphanes camena* from the combined morphological and molecular dataset (total-evidence) because its phylogenetic position in the phylogeny was unstable (Fig. S2B, G). We selected one specimen per species to ensure a species-level phylogeny. The concatenated total-evidence analysis was run in MrBayes v.3.2.6 via the CIPRES. We used the best-fit partitioning strategy suggested by PartitionFinder v.2.1.1 for the molecular data (Table S2). We partitioned the morphological data by using homoplasy scores calculated through implied weighting parsimony (Table S3), as detailed in the main text and in (Rosa *et al.*, 2019). The analysis was set as above ('*Concatenated molecular data*') and using the Markov (MKv) model for the morphological data (Lewis, 2001). We summarized the post-burnin trees using the 50% majority-rule consensus method (Fig. S3).

*Total-evidence species tree using the multispecies coalescent model*

Molecular clock test

We compared the strict and relaxed clocks (Thorne & Kishino, 2002) for the molecular dataset using Bayes factors (Kass & Raftery, 1995). We ran stepping-stone sampling analyses in MrBayes v.3.2.6 for 50 million generations, sampling every 5,000 generations. Marginal likelihoods were used to compute twice the natural logarithm of the Bayes factors (2 $\log_e$ *BF*), and we considered values > 10 to provide very strong evidence against the strict clock model. The relaxed clock model was then preferred for all loci (Table S4).

Tree model test

We evaluated the fit of two tree models available in StarBEAST2 v.0.15.5 (Ogilvie *et al.*, 2017): the Yule and birth-death models. In addition, we evaluated the fit of three molecular clock partitions: one single clock, two clocks (mitochondrial and nuclear), and six clocks (one for each gene partition). The analyses were set as described in the main text using BEAST v.2.6.3 (Bouckaert *et al.*, 2014). We set 25 path-sampling steps using thermodynamic integration (Lartillot & Philippe, 2006), each step running for 60 million generations. We evaluated convergence by checking that the estimated sample sizes (ESS) were above 200 in every path-sampling step. The marginal likelihood estimate for the Yule tree model and the two-clock partition was −64584.43, and was decisively supported for the Brassolini dataset based on Bayes factor comparisons (Table S5).

Gene-tree discordance test

We estimated the contribution of each gene tree in a multi-locus phylogenetic analysis using partitioned Bremer support scores (Baker & DeSalle, 1997). The analysis was carried out in TNT v.1.5 (Goloboff & Catalano, 2016) and using a script written by Peña *et al.* (2006). Although the phylogenetic signal is low to moderate in the nodes close to the root, there are conflicts among gene partitions. For example, the Brassolini clade excluding the genus *Bia* received disparate support: COI, 0.6; RpS5, −0.5; GAPDH, 8.5; EF1α, −8.6; CAD, 0; *wingless*, 3.0. This might indicate that the low posterior probabilities close to the root of Brassolini may be related to gene tree conflict (Fig. S4); thus, acknowledging such discordances via the multispecies coalescent might alleviate potential biases in species tree topology and divergence time inference.

Species tree topology test

We evaluated the likelihood of the branching orders among early divergent Brassolini lineages. Specifically, we assessed the main discrepancy among 1) the morphology-based systematics of Brassolini (Penz, 2007), 2) the total-evidence consensus phylogeny using the concatenation approach (Fig. S3), and 3) the multispecies coalescent MCC species tree (Fig. S5). That is, respectively, 1) *Narope* (former subtribe Naropina) sister to the remaining Brassolina (i.e., all Brassolini genera but the genus *Bia*), 2) *Brassolis* sister to the *Opsiphanes*-group (Clade D in Fig. 2), and 3) *Brassolis* sister to the *Opsiphanes*- and *Caligo*-groups (Clades C + D in Fig. 2).

We inferred three maximum-likelihood molecular phylogenies with the nodes under investigation constrained using IQ-TREE v.2.0.5 (Minh *et al.*, 2020). The multi-locus dataset was partitioned as suggested by PartitionFinder v.2.1.1 (Table S2) and we let ModelFinder (Kalyaanamoorthy *et al.*, 2017) implemented in IQ-TREE v2.0.5 select the best-fit substitution models. We carried out tree topology tests by 1) approximating bootstrap proportions by resampling 10,000 times the estimated log-likelihoods of sites (Kishino *et al.*, 1990), 2) estimating expected likelihood weights (Strimmer & Rambaut, 2002), 3) performing weighted KH (Kishino & Hasegawa, 1989) and SH tests (Shimodaira & Hasegawa, 1999), and 4) carrying out the approximately unbiased (AU) test (Shimodaira, 2002). The tree topology tests did not reject either branching order inferred by the concatenation or the multi-species coalescent approaches (Table S6), suggesting that the molecular dataset is in agreement with both total-evidence consensus and MCC species trees (Figs S3 and S5). The tree topology depicting *Narope* as sister to Brassolina is clearly rejected; thus, Naropina is subsumed within Brassolina.

*Taxonomic resolution across Neotropical bioregions*

To assess any biases in macroevolutionary analyses due to disparate taxonomic effort across the Neotropics, we compiled the year of description and species revisions of every Brassolini species per bioregion from (Lamas, 2004; Austin *et al.*, 2007; Bristow, 2008; Penz, 2008, 2009; Garzón-Orduña & Penz, 2009; Penz *et al.*, 2011; Penz *et al.*, 2017; Chacón *et al.*, 2012).

First, we evaluated regional rates of species descriptions which might depict potential geographical biases in collecting and describing taxa. We found that, from the mid-XIX century to the first quarter of the XX century, there has been a vivid taxonomic activity across all Neotropical bioregions. This resulted in the collection and description of ~80% of endemic species in Mesoamerica, Amazonia, and the Atlantic Forest by 1925 (Fig. S6A, C).

Second, we evaluated regional rates of taxonomical revisions which might depict potential biases in oversplitting species in a particular biome. There has been a recent increase of studies describing/revising species across all Neotropical bioregions (Fig. S6D, F), though the trend is less pronounced in the Atlantic Forest. This might not have been driven by either a conservative criterion of taxonomists working in such a region or an oversplitting criterion of taxonomists working in Mesoamerica and Amazonia, because most described/revised species during the past 35 years come from genus-level studies using specimens from across the Neotropics, mainly by M. Casagrande (revision of *Narope* 1989, 2002), C. Penz, G. Austin, I. Garzón-Orduña, P. DeVries and colleagues (revisions of *Bia*, *Blepolenis*, *Brassolis*, *Dasyophthalma*, *Dynastor*, *Eryphanis*, *Opoptera*, *Opsiphanes*, *Orobrassolis*).

Third, we assessed any geographical biases in describing multiple infraspecific taxa which might indicate a tendency for lumping species. In particular, we evaluated whether the recent increase in speciation rate in Mesoamerica was driven by a trend to treat subspecies as full species only in such a region. From the 36 Brassolini species occurring in more than one Neotropical bioregion, 28 species have multiple allopatric infraspefic taxa (i.e., subspecies). From these, the number of polytypic species occurring in Mesoamerica is slightly higher than Atlantic Forest polytypic species (11 *vs*. 7 species, respectively). Therefore, it seems that there is no tendency to treat subspecies as full species in Mesoamerica which otherwise would have inflated speciation rates.

Fourth, we evaluated whether our taxon sampling has been biased towards a particular region. From 108 described Brassolini species, 72 species (67%) occur in only one of the defined areas, and from these, 33 species (31%) occur on either Mesoamerica or the Atlantic Forest. In our study using molecular and/or morphological characters, from the 84 examined Brassolini species, 49 species (58%) occur in only one of the defined areas, and from these, 28 species (33%) occur on either Mesoamerica or the Atlantic Forest. This suggests that our incomplete sampling reflects the actual biogeographical signal of extant Brassolini species.

*Missing species*

We aimed to take into account the 25 unsampled species in our phylogeny for the estimation of dispersal and speciation rates. In the program BioGeoBEARS, we randomly added missing lineages to their currently assigned monophyletic genera in 100 posterior species trees. In the program BAMM, we generated clade-specific sampling fractions at the genus level. The proportions of missing taxa in both analyses follow Table S7. Note that we also repeated the analyses using the inferred species tree without considering missing taxa (Figs S7, S8, and S9).

### Figures & Tables

*Figure S1*: Map of the Neotropics showing the 7,174 cleaned occurrences of Brassolini species and the 81 flagged as potential errors, which were excluded from the bioregion delimitation analyses.

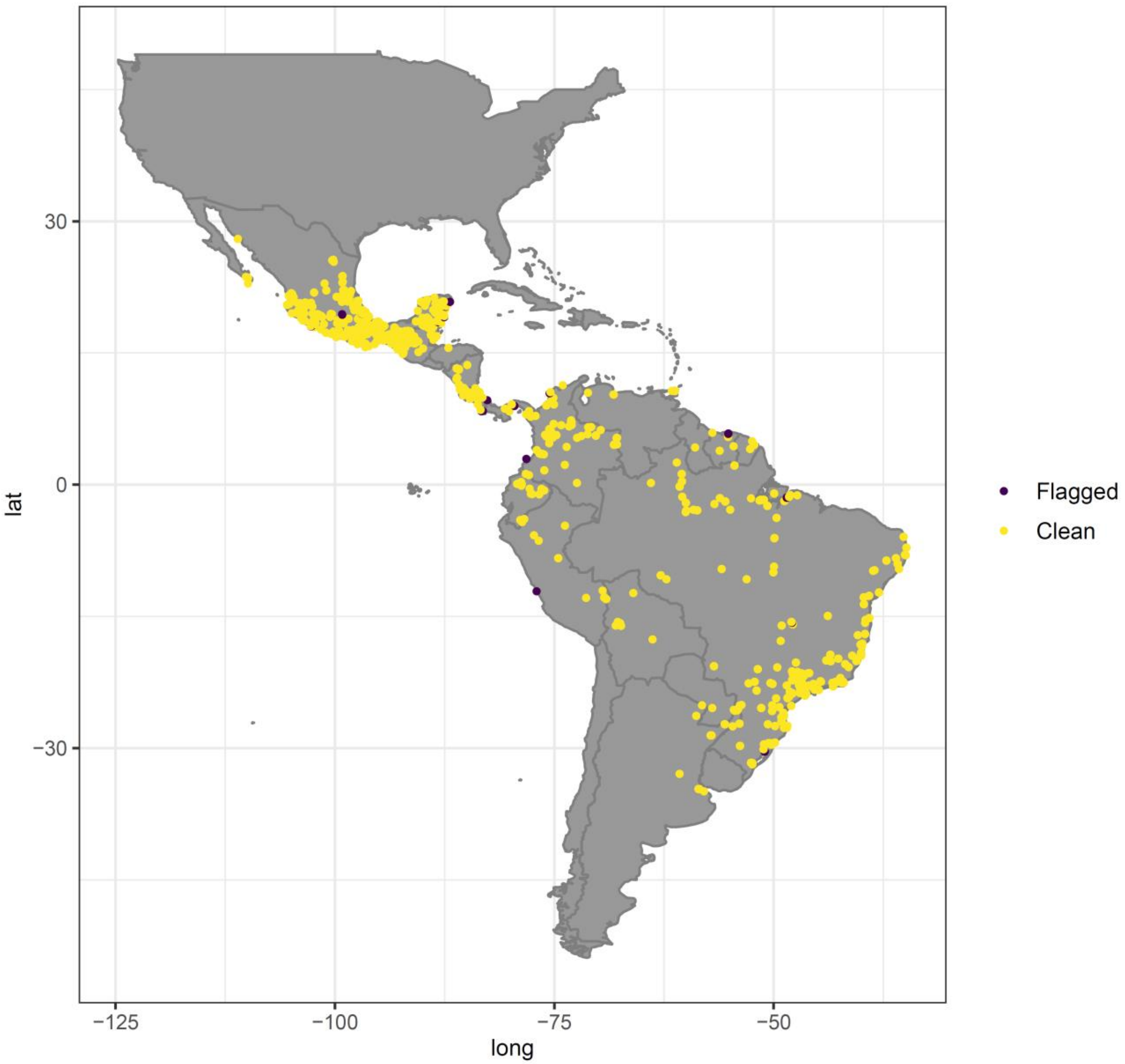

*Figure S2*: Single-gene, concatenated multi-locus and morphological trees. Each tree represents the 50% majority-rule consensus of 7,500 posterior trees inferred in MrBayes v.3.2.6. Posterior probabilities are shown on every node. A: CAD gene tree; B: COI gene tree; C: EF1α gene tree; D: GAPDH gene tree; E: RpS5 gene tree; F: *wingless* gene tree; G: concatenated multi-locus molecular tree; H: morphology-based tree.

A: CAD gene tree

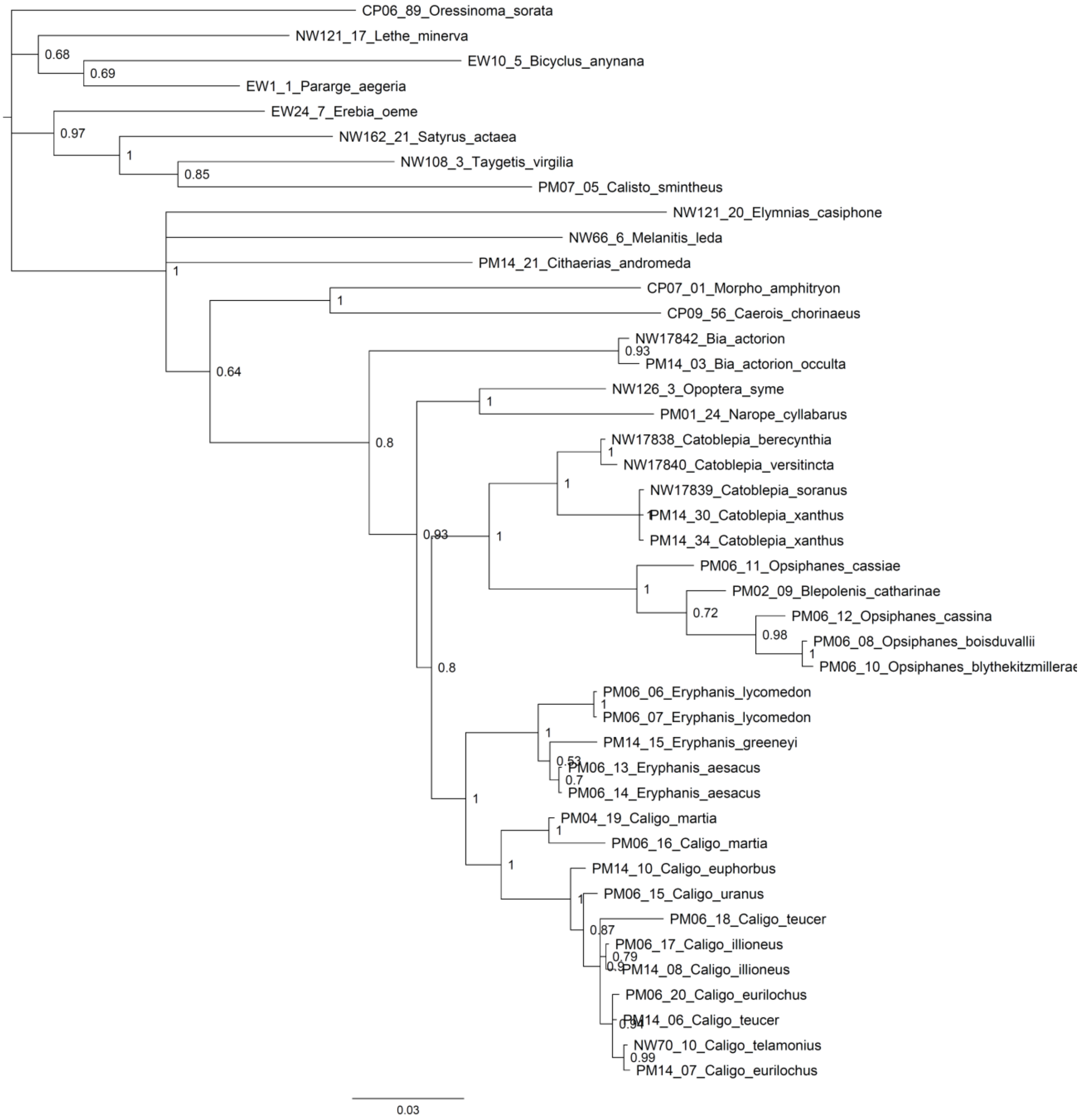

B: COI gene tree

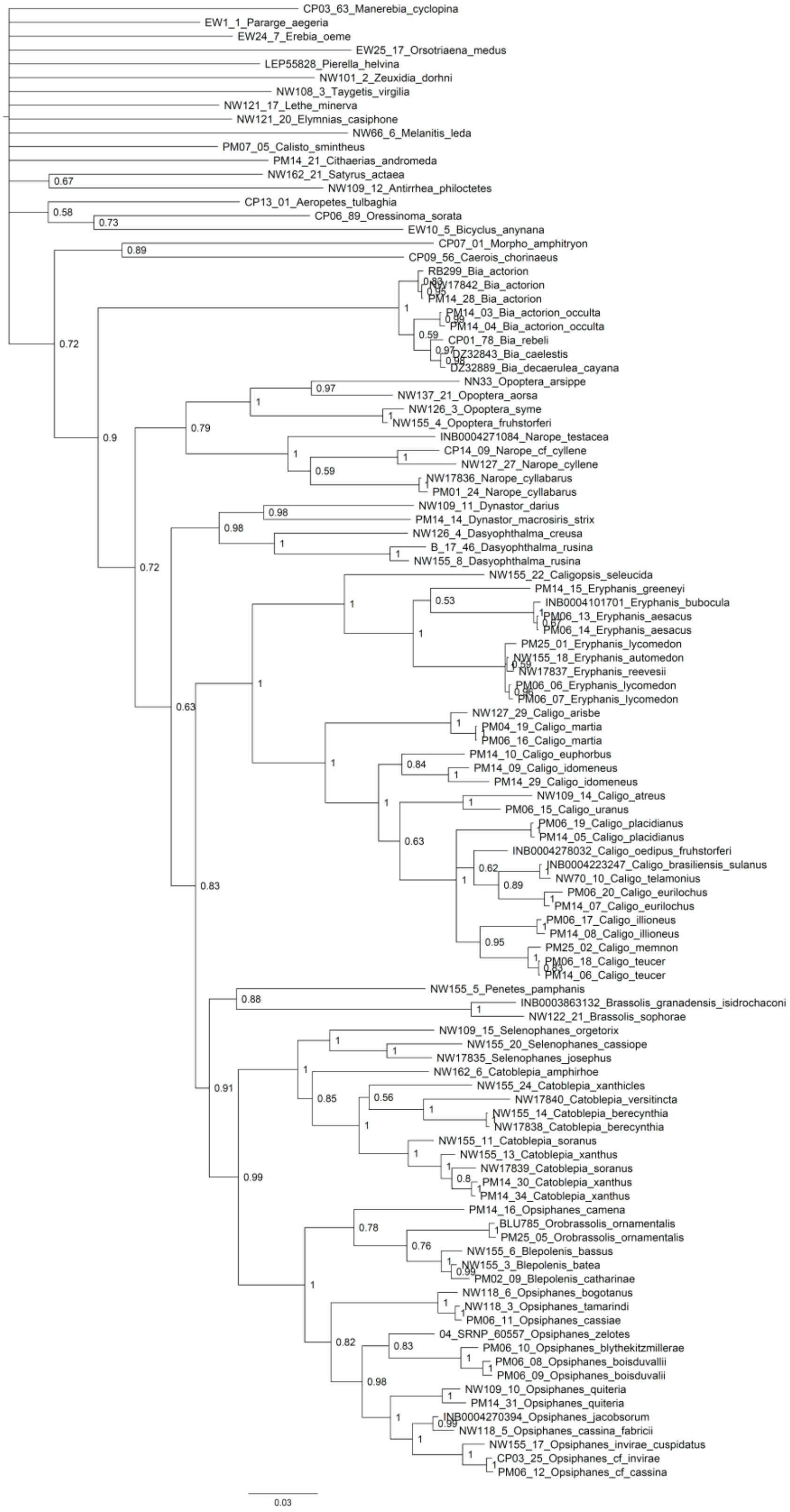

C: EF1α gene tree

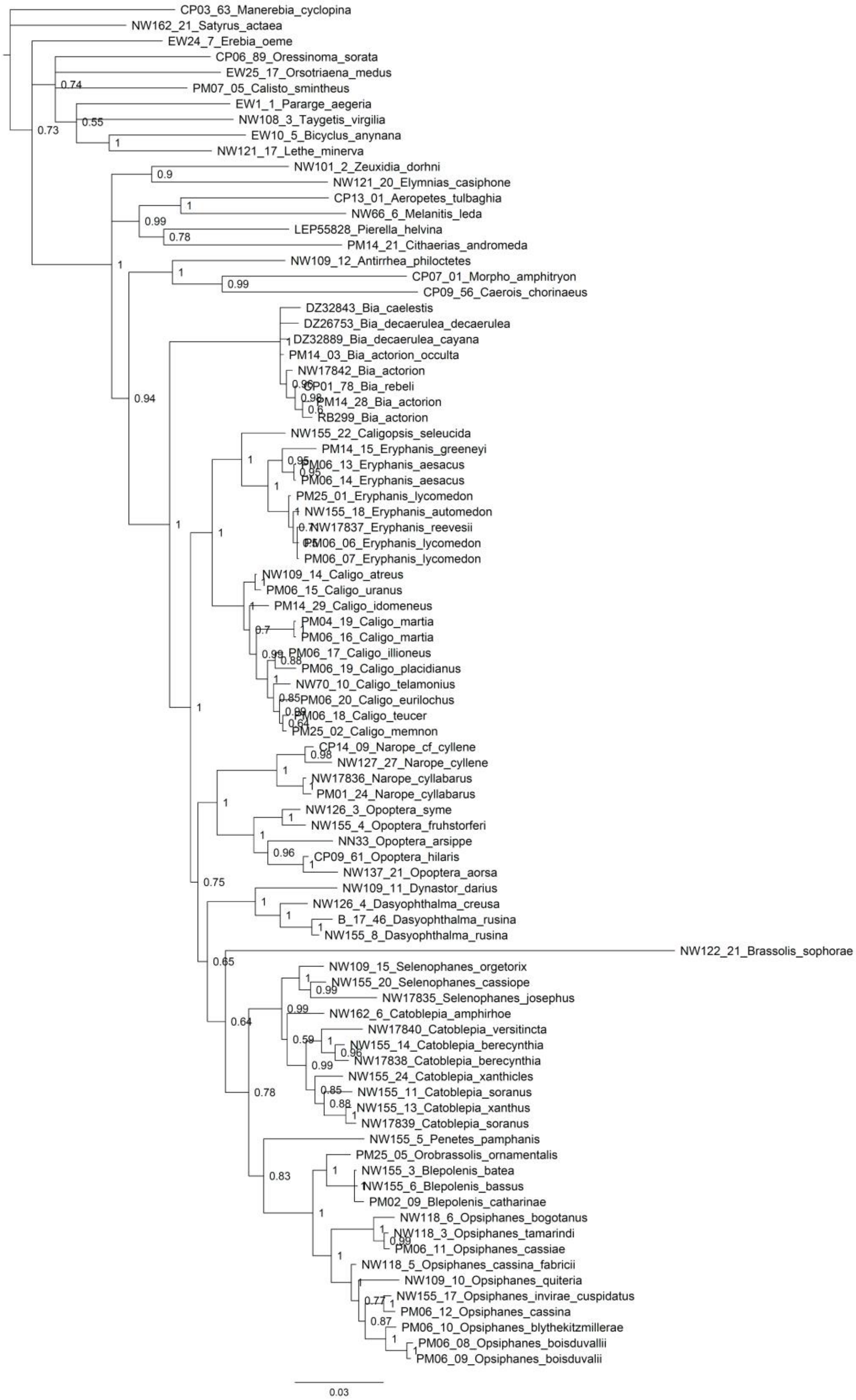

D: GAPDH gene tree

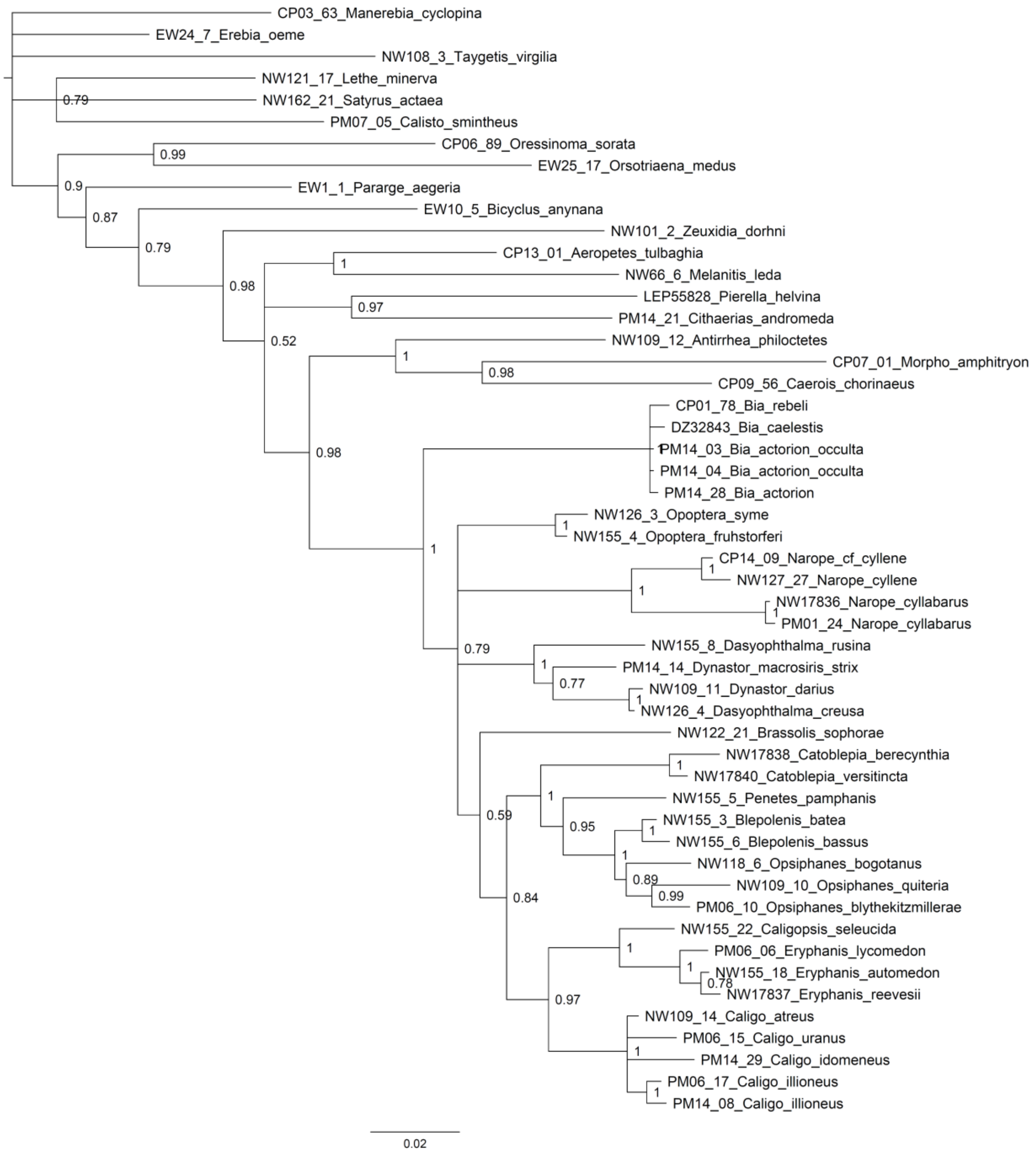

E: RpS5 gene tree

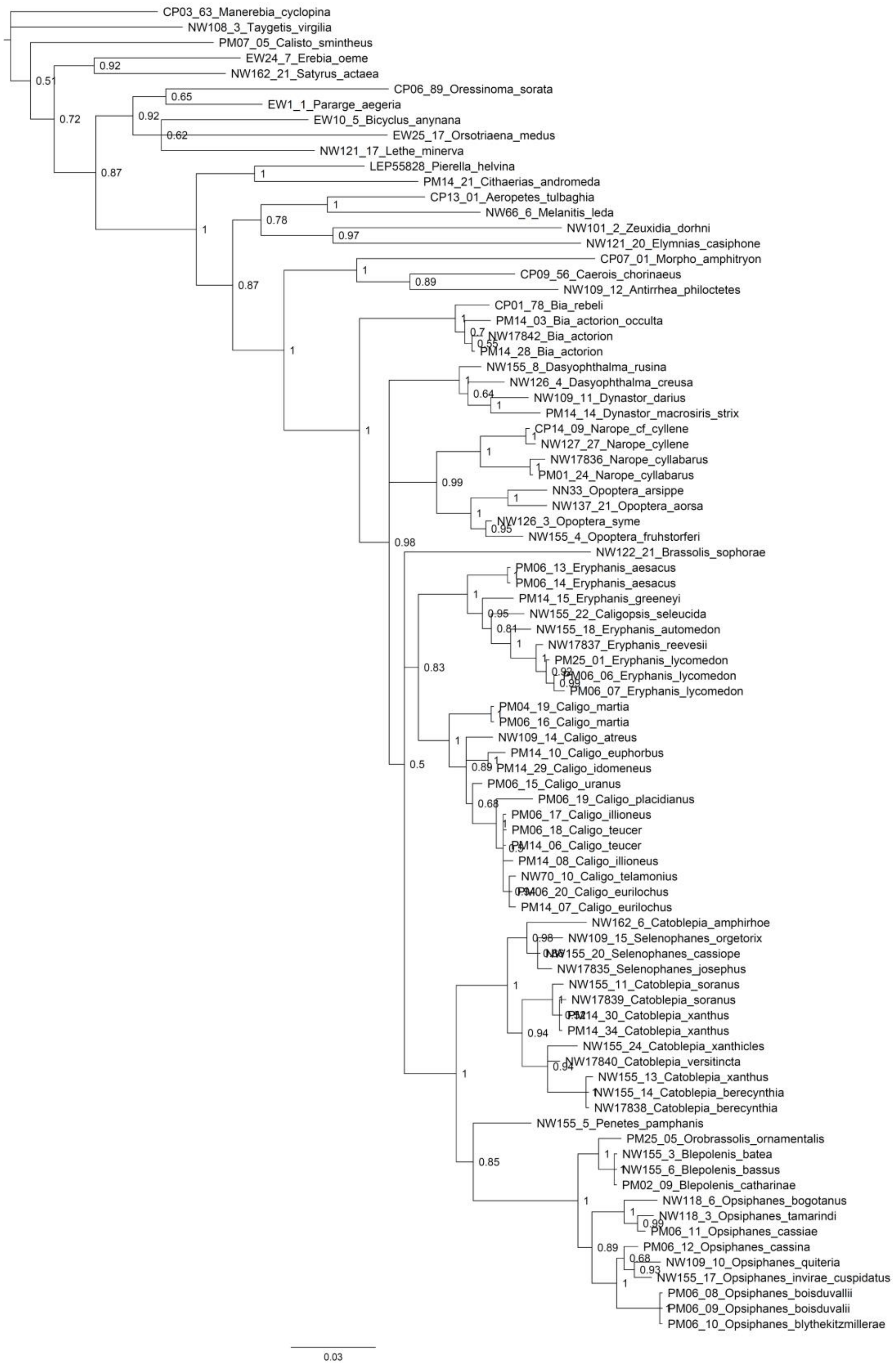

F: *wingless* gene tree

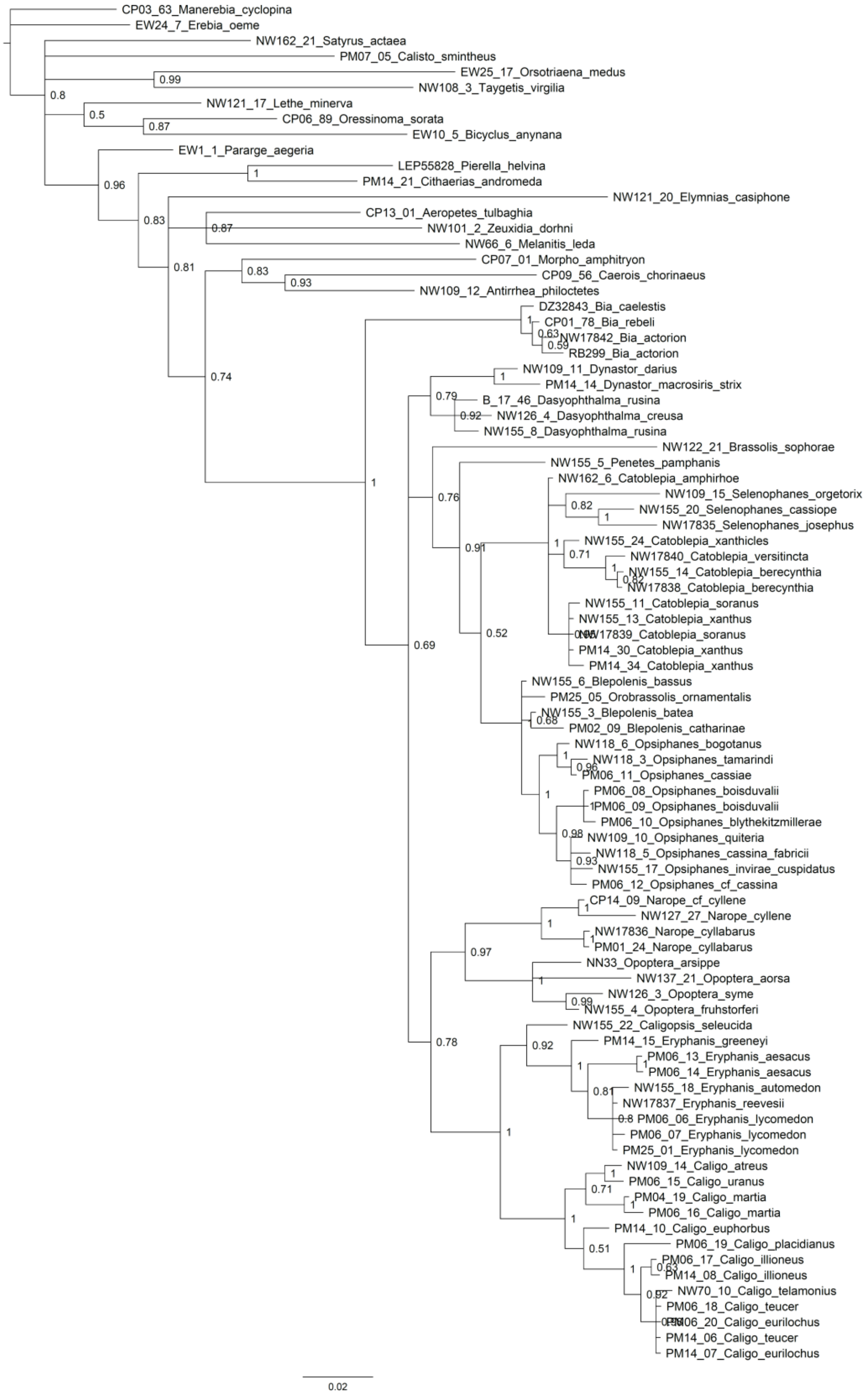

G: concatenated multi-locus molecular tree.

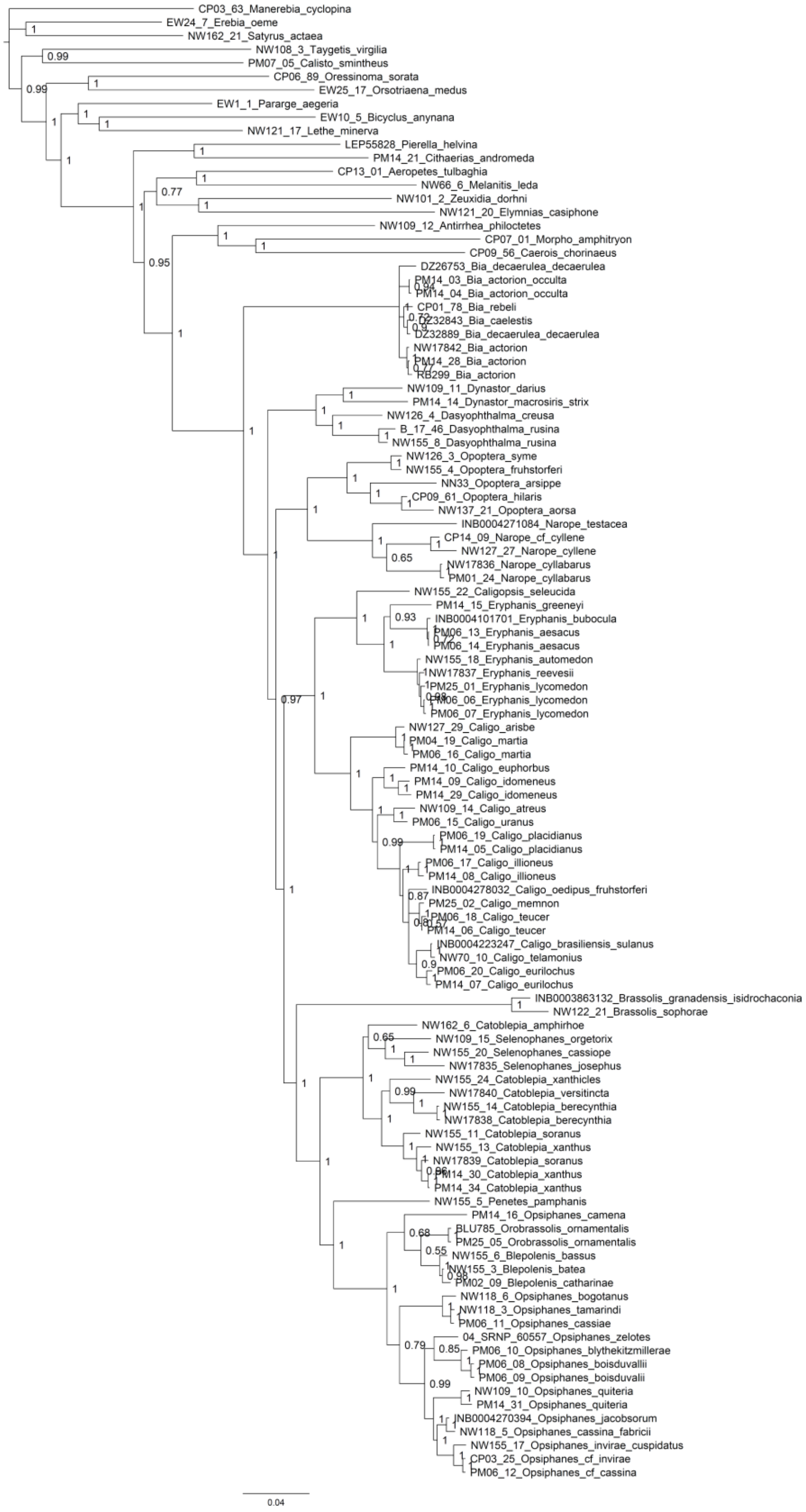

H: morphology-based tree

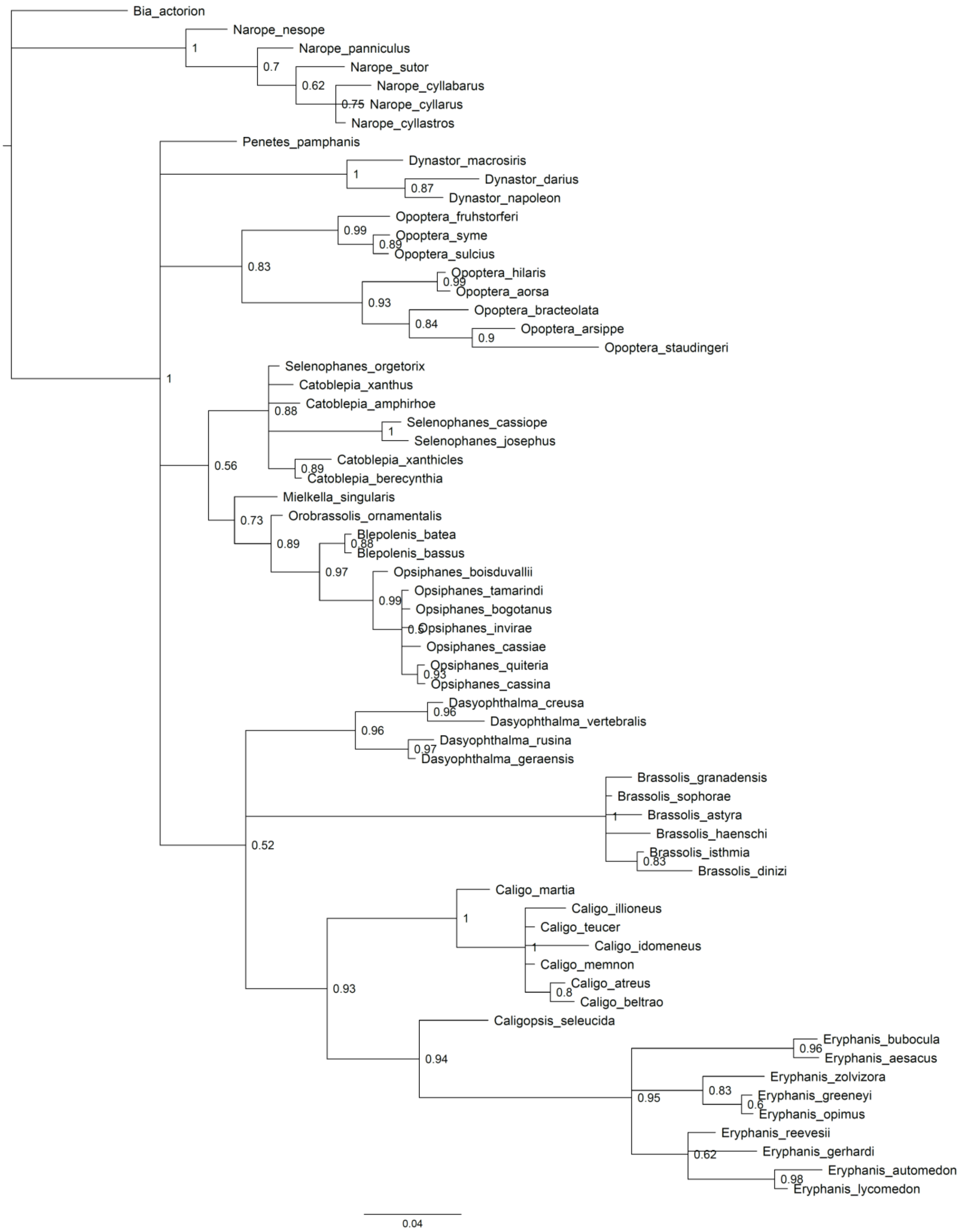

*Figure S3*: Total-evidence consensus tree using the concatenation approach. The tree represents the 50% majority-rule consensus of 7,500 posterior trees inferred in MrBayes v.3.2.6. Posterior probabilities are shown on every node.

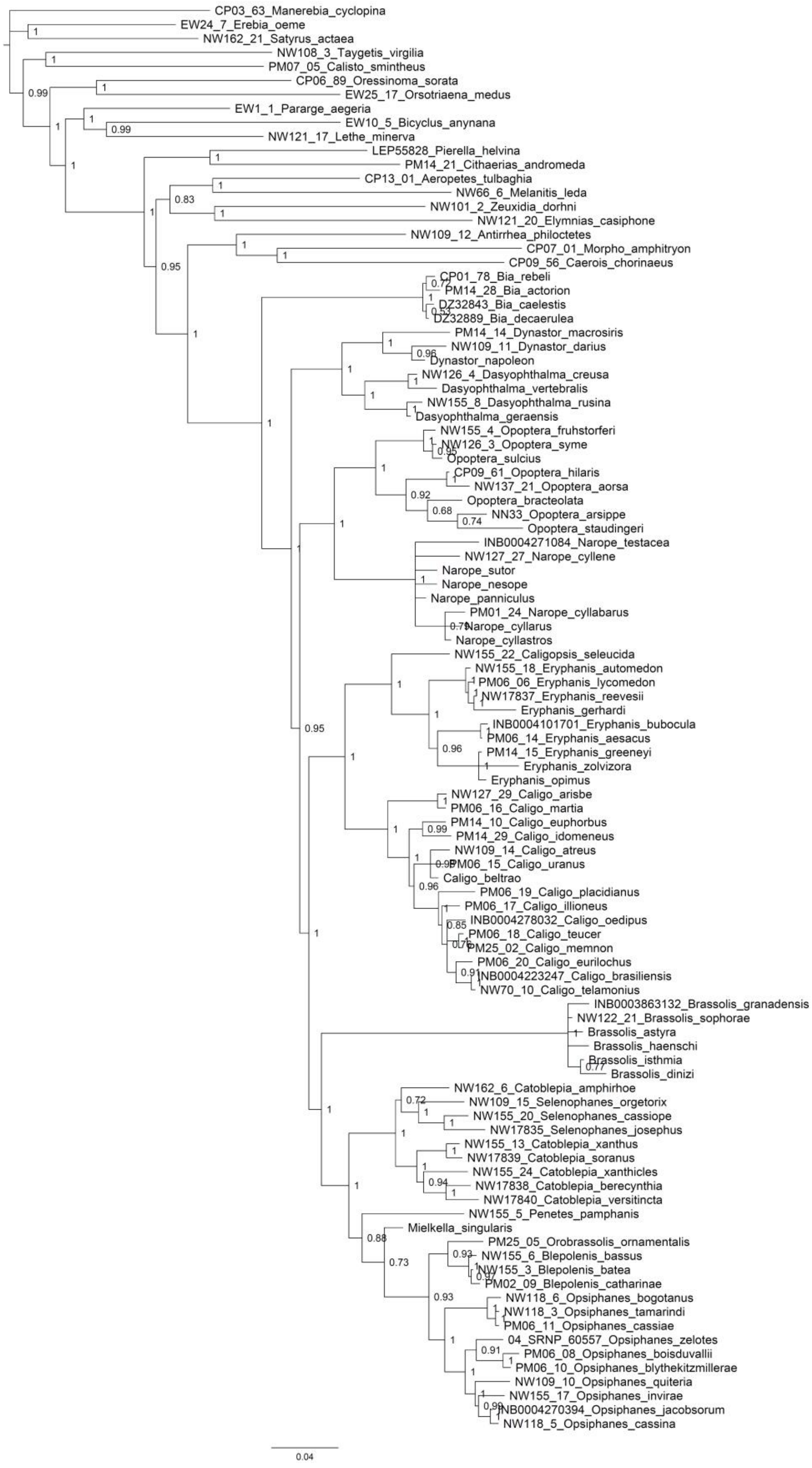
CP03_63_Manerebia_cyclopina
EW24_7_Erebia_oeme
NW162_21_Satyrus_actaea
NW108_3_Taygetis_virgilia
PM07_05_Calisto_smintheus
CP06_89_Oressinoma_sorata
EW25_17_Orsotriaena_medus
EW1_1_Pararge_aegeria
EW10_5_Bicyclus_anynana
NW121_17_Lethe_minerva
LEP55828_Pierella_helvina
PM14_21_Cithaerias_andromeda
CP13_01_Aeropetes_tulbaghia
NW66_6_Melanitis_leda
NW101_2_Zeuxidia_dorhni
NW121_20_Elymnias_casiphone
NW109_12_Antirrhea_philoctetes
CP07_01_Morpho_amphitryon
CP09_56_Caerois_chorinaeus
CP01_78_Bia_rebeli
PM14_28_Bia_actorion
DZ32843_Bia_caelestis
DZ32889_Bia_decaerulea
PM14_14_Dynastor_macrosiris
NW109_11_Dynastor_darius
Dynastor_napoleon
NW126_4_Dasyophthalma_creusa
Dasyophthalma_vertebralis
NW155_8_Dasyophthalma_rusina
Dasyophthalma_geraensis
NW155_4_Opoptera_fruhstorferi
NW126_3_Opoptera_syme
Opoptera_sulcius
CP09_61_Opoptera_hilaris
NW137_21_Opoptera_aorsa
Opoptera_bracteolata
NN33_Opoptera_arsippe
Opoptera_staudingeri
INB0004271084_Narope_testacea
NW127_27_Narope_cyllene
Narope_sutor
Narope_nesope
Narope_panniculus
PM01_24_Narope_cyllabarus
Narope_cyllarus
Narope_cyllastros
NW155_22_Caligopsis_seleucida
NW155_18_Eryphanis_automedon
PM06_06_Eryphanis_lycomedon
NW17837_Eryphanis_reevesii
Eryphanis_gerhardi
INB0004101701_Eryphanis_bubocula
PM06_14_Eryphanis_aesacus
PM14_15_Eryphanis_greeneyi
Eryphanis_zolvizora
Eryphanis_opimus
NW127_29_Caligo_arisbe
PM06_16_Caligo_martia
PM14_10_Caligo_euphorbus
PM14_29_Caligo_idomeneus
NW109_14_Caligo_atreus
PM06_15_Caligo_uranus
Caligo_beltrao
PM06_19_Caligo_placidianus
PM06_17_Caligo_illioneus
INB0004278032_Caligo_oedipus
PM06_18_Caligo_teucer
PM25_02_Caligo_memnon
PM06_20_Caligo_eurilochus
INB0004223247_Caligo_brasiliensis
NW70_10_Caligo_telamonius
INB0003863132_Brassolis_granadensis
NW122_21_Brassolis_sophorae
Brassolis_astyra
Brassolis_haenschi
Brassolis_isthmia
Brassolis_dinizi
NW162_6_Catoblepia_amphirhoe
NW109_15_Selenophanes_orgetorix
NW155_20_Selenophanes_cassiope
NW17835_Selenophanes_josephus
NW155_13_Catoblepia_xanthus
NW17839_Catoblepia_soranus
NW155_24_Catoblepia_xanthicles
NW17838_Catoblepia_berecynthia
NW17840_Catoblepia_versitincta
NW155_5_Penetes_pamphanis
Mielkella_singularis
PM25_05_Orobrassolis_ornamentalis
NW155_6_Blepolenis_bassus
NW155_3_Blepolenis_batea
PM02_09_Blepolenis_catharinae
NW118_6_Opsiphanes_bogotanus
NW118_3_Opsiphanes_tamarindi
PM06_11_Opsiphanes_cassiae
04_SRNP_60557_Opsiphanes_zelotes
PM06_08_Opsiphanes_boisduvallii
PM06_10_Opsiphanes_blythekitzmillerae
NW109_10_Opsiphanes_quiteria
NW155_17_Opsiphanes_invirae
INB0004270394_Opsiphanes_jacobsorum
NW118_5_Opsiphanes_cassina
0.04

*Figure S4*: Strict consensus tree of the 13 equally parsimonious trees estimated using maximum parsimony in TNT v.1.5. Numbers next to nodes represent the contribution of genes as measured by partitioned Bremer support. The scores correspond to the genes CAD, COI, EF1α, GAPDH, RpS5, and *wingless*, respectively.

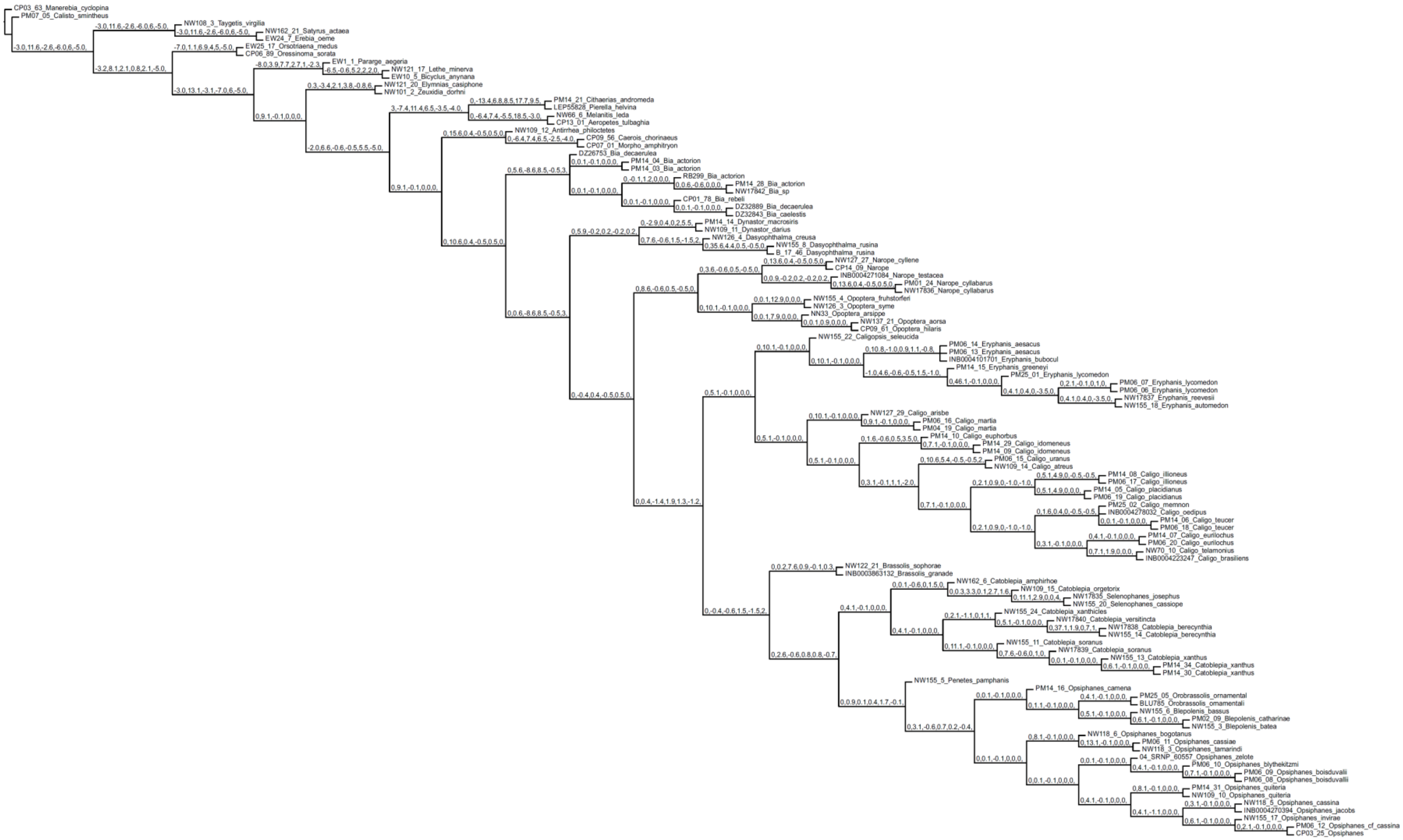
CP03_63_Manerebia_cyclopina
PM07_05_Calisto_smintheus
NW108_3_Taygetis_virgilia
NW162_21_Satyrus_actaea
EW24_7_Erebia_oeme
EW25_17_Orsotriaena_medus
CP06_89_Oressinoma_sorata
EW1_1_Pararge_aegeria
NW121_17_Lethe_minerva
EW10_5_Bicyclus_anynana
NW121_20_Elymnias_casiphone
NW101_2_Zeuxidia_dorhni
PM14_21_Cithaerias_andromeda
LEP55828_Pierella_helvina
NW66_6_Melanitis_leda
CP13_01_Aeropetes_tulbaghia
NW109_12_Antirrhea_philoctetes
CP09_56_Caerois_chorinaeus
CP07_01_Morpho_amphitryon
DZ26753_Bia_decaerulea
PM14_04_Bia_actorion
PM14_03_Bia_actorion
RB299_Bia_actorion
PM14_28_Bia_actorion
NW17842_Bia_sp
CP01_78_Bia_rebeli
DZ32889_Bia_decaerulea
DZ32843_Bia_caelestis
PM14_14_Dynastor_macrosiris
NW109_11_Dynastor_darius
NW126_4_Dasyophthalma_creusa
NW155_8_Dasyophthalma_rusina
B_17_46_Dasyophthalma_rusina
NW127_27_Narope_cyllene
CP14_09_Narope
INB0004271084_Narope_testacea
PM01_24_Narope_cyllabarus
NW17836_Narope_cyllabarus
NW155_4_Opoptera_fruhstorferi
NW126_3_Opoptera_syme
NN33_Opoptera_arsippe
NW137_21_Opoptera_aorsa
CP09_61_Opoptera_hilaris
NW155_22_Caligopsis_seleucida
PM06_14_Eryphanis_aesacus
PM06_13_Eryphanis_aesacus
INB0004101701_Eryphanis_bubocul
PM14_15_Eryphanis_greeneyi
PM25_01_Eryphanis_lycomedon
PM06_07_Eryphanis_lycomedon
PM06_06_Eryphanis_lycomedon
NW17837_Eryphanis_reevesii
NW155_18_Eryphanis_automedon
NW127_29_Caligo_arisbe
PM06_16_Caligo_martia
PM04_19_Caligo_martia
PM14_10_Caligo_euphorbus
PM14_29_Caligo_idomeneus
PM14_09_Caligo_idomeneus
PM06_15_Caligo_uranus
NW109_14_Caligo_atreus
PM14_08_Caligo_illioneus
PM06_17_Caligo_illioneus
PM14_05_Caligo_placidianus
PM06_19_Caligo_placidianus
PM25_02_Caligo_memnon
INB0004278032_Caligo_oedipus
PM14_06_Caligo_teucer
PM06_18_Caligo_teucer
PM14_07_Caligo_eurilochus
PM06_20_Caligo_eurilochus
NW70_10_Caligo_telamonius
INB0004223247_Caligo_brasiliens
NW122_21_Brassolis_sophorae
INB0003863132_Brassolis_granade
NW162_6_Catoblepia_amphirhoe
NW109_15_Catoblepia_orgetorix
NW17835_Selenophanes_josephus
NW155_20_Selenophanes_cassiope
NW155_24_Catoblepia_xanthicles
NW17840_Catoblepia_versitincta
NW17838_Catoblepia_berecynthia
NW155_14_Catoblepia_berecynthia
NW155_11_Catoblepia_soranus
NW17839_Catoblepia_soranus
NW155_13_Catoblepia_xanthus
PM14_34_Catoblepia_xanthus
PM14_30_Catoblepia_xanthus
NW155_5_Penetes_pamphanis
PM14_16_Opsiphanes_camena
PM25_05_Orobrassolis_ornamental
BLU785_Orobrassolis_ornamentali
NW155_6_Blepolenis_bassus
PM02_09_Blepolenis_catharinae
NW155_3_Blepolenis_batea
NW118_6_Opsiphanes_bogotanus
PM06_11_Opsiphanes_cassiae
NW118_3_Opsiphanes_tamarindi
04_SRNP_60557_Opsiphanes_zelote
PM06_10_Opsiphanes_blythekitzmi
PM06_09_Opsiphanes_boisduvallii
PM06_08_Opsiphanes_boisduvallii
PM14_31_Opsiphanes_quiteria
NW109_10_Opsiphanes_quiteria
NW118_5_Opsiphanes_cassina
INB0004270394_Opsiphanes_jacobs
NW155_17_Opsiphanes_invirae
PM06_12_Opsiphanes_cf_cassina
CP03_25_Opsiphanes

*Figure S5*: Maximum clade credibility species tree using the total-evidence dataset and the multispecies coalescent model in BEAST v.2.6.3. Posterior probabilities and the 95% HPD (highest posterior density) intervals are shown on every node. The species tree is calibrated in million years. Calibration points based on Chazot *et al.*, 2019a, are indicated by red arrows: 1) Satyrinae (46–65 Mya), 2) divergence Melanitini and Dirini (27–41 Mya), 3) Satyrini (38–55 Mya), 4) crown node of Lethina, Parargina and Mycalesina (31–46 Mya), 5) crown node of Pronophilina, Euptychiina, Satyrina and Erebiina (31–44 Mya), 6) divergence Brassolini and Morphini (34–50 Mya).

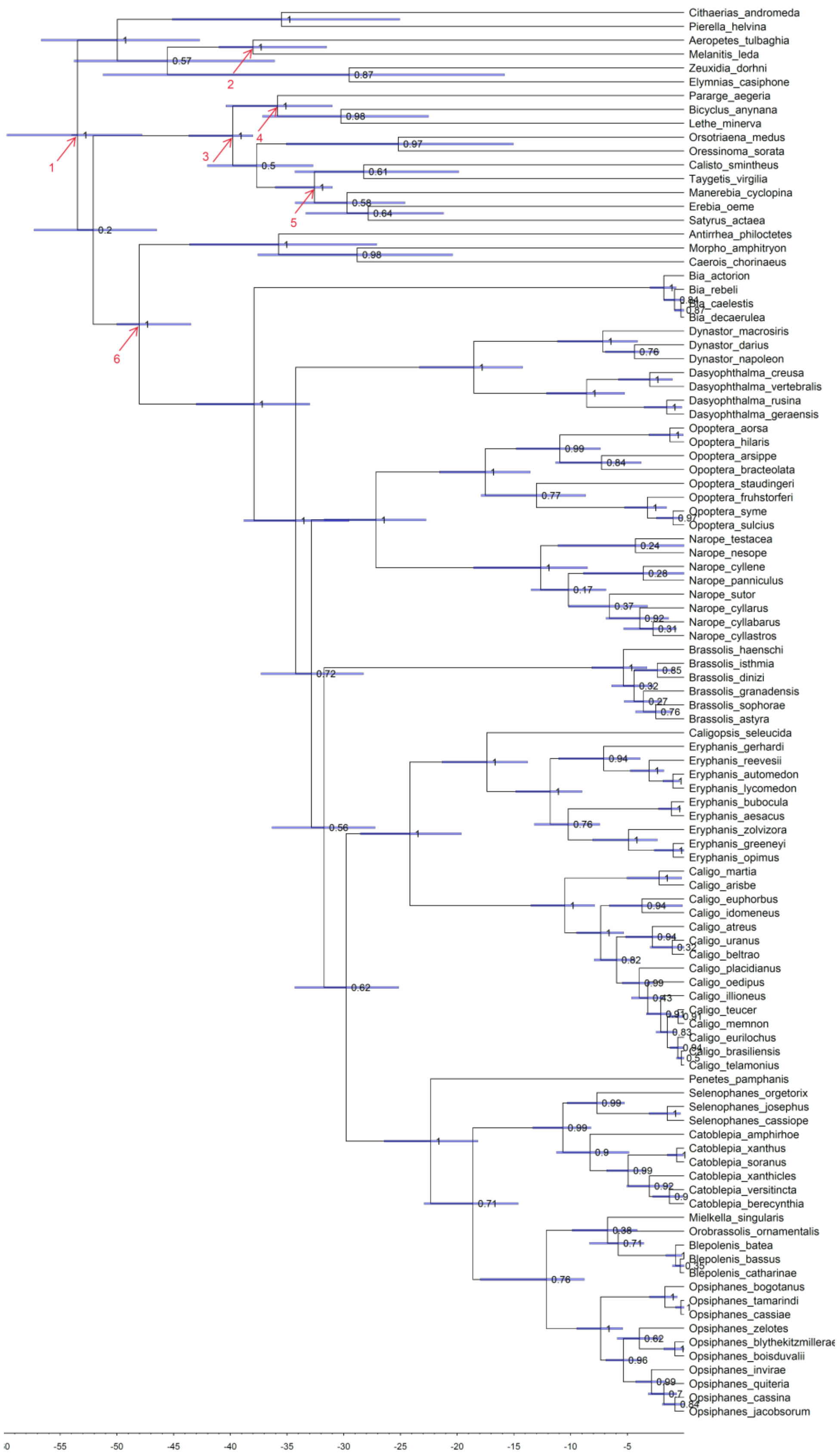

Cithaerias_andromeda
Pierella_helvina
Aeropetes_tulbaghia
Melanitis_leda
Zeuxidia_dorhni
Elymnias_casiphone
Pararge_aegeria
Bicyclus_anynana
Lethe_minerva
Orsotriaena_medus
Oressinoma_sorata
Calisto_smintheus
Taygetis_virgilia
Manerebia_cyclopina
Erebia_oeme
Satyrus_actaea
Antirrhea_philoctetes
Morpho_amphitryon
Caerois_chorinaeus
Bia_actorion
Bia_rebeli
Bia_caelestis
Bia_decaerulea
Dynastor_macrosiris
Dynastor_darius
Dynastor_napoleon
Dasyophthalma_creusa
Dasyophthalma_vertebralis
Dasyophthalma_rusina
Dasyophthalma_geraensis
Opoptera_aorsa
Opoptera_hilaris
Opoptera_arsippe
Opoptera_bracteolata
Opoptera_staudingeri
Opoptera_fruhstorferi
Opoptera_syme
Opoptera_sulcius
Narope_testacea
Narope_nesope
Narope_cyllene
Narope_panniculus
Narope_sutor
Narope_cyllarus
Narope_cyllabarus
Narope_cyllastros
Brassolis_haenschi
Brassolis_isthmia
Brassolis_dinizi
Brassolis_granadensis
Brassolis_sophorae
Brassolis_astyra
Caligopsis_seleucida
Eryphanis_gerhardi
Eryphanis_reevesii
Eryphanis_automedon
Eryphanis_lycomedon
Eryphanis_bubocula
Eryphanis_aesacus
Eryphanis_zolvizora
Eryphanis_greeneyi
Eryphanis_opimus
Caligo_martia
Caligo_arisbe
Caligo_euphorbus
Caligo_idomeneus
Caligo_atreus
Caligo_uranus
Caligo_beltrao
Caligo_placidianus
Caligo_oedipus
Caligo_illioneus
Caligo_teucer
Caligo_memnon
Caligo_eurilochus
Caligo_brasiliensis
Caligo_telamonius
Penetes_pamphanis
Selenophanes_orgetorix
Selenophanes_josephus
Selenophanes_cassiope
Catoblepia_amphirhoe
Catoblepia_xanthus
Catoblepia_soranus
Catoblepia_xanthicles
Catoblepia_versitincta
Catoblepia_berecynthia
Mielkella_singularis
Orobrassolis_ornamentalis
Blepolenis_batea
Blepolenis_bassus
Blepolenis_catharinae
Opsiphanes_bogotanus
Opsiphanes_tamarindi
Opsiphanes_cassiae
Opsiphanes_zelotes
Opsiphanes_blythekitzmillerae
Opsiphanes_boisduvalii
Opsiphanes_invirae
Opsiphanes_quiteria
Opsiphanes_cassina
Opsiphanes_jacobsorum
1
2
3
4
5
6
-55
-50
-45
-40
-35
-30
-25
-20
-15
-10
-5


9

*Figure S6*: Brassolini taxonomic resolution across Neotropical bioregions. We found similar trends across Mesoamerica, Amazonia and Atlantic Forest in collecting, describing and revising the species status of all valid Brassolini taxa. Thus, we rule out any taxonomic bias affecting our biogeographical and diversification analyses. First, we compiled the year of description of every species restricted to A) Amazonia, B) Mesoamerica and NW Andes, and C) Atlantic Forest. Second, we compiled the year of the last revision of the specific status of Brassolini taxa restricted to D) Amazonia, E) Mesoamerica and NW Andes, and C) Atlantic Forest. Vivid taxonomic activity has occurred simultaneously in the three main rainforest biomes in the Neotropics during the mid-XIX century to the first quarter of the XX century, and during the late XX century to the present.

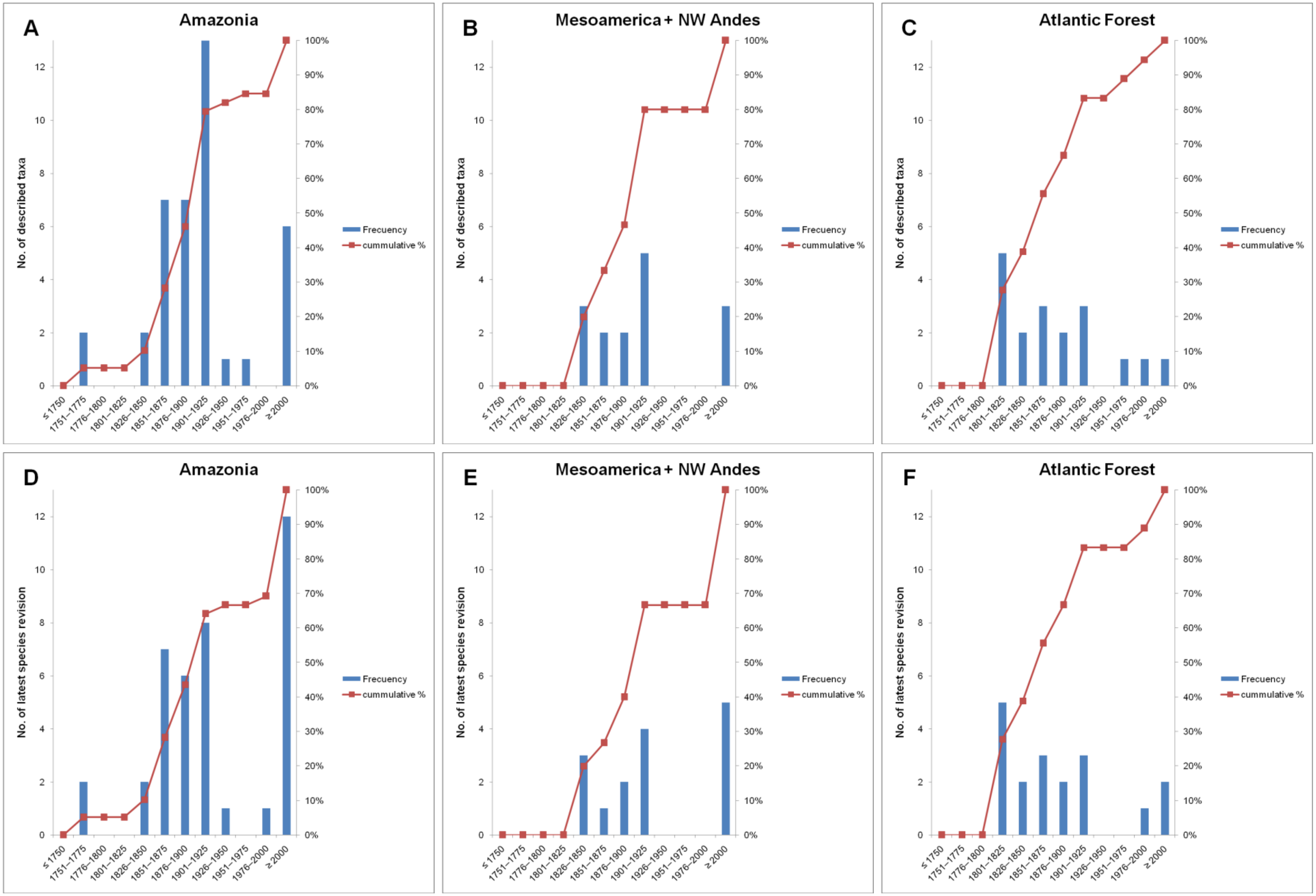
A
Amazonia
B
Mesoamerica + NW Andes
C
Atlantic Forest
D
Amazonia
E
Mesoamerica + NW Andes
F
Atlantic Forest
No. of described taxa
No. of latest species revision
Frecuency
cummulative %
≤ 1750
1751–1775
1776–1800
1801–1825
1826–1850
1851–1875
1876–1900
1901–1925
1926–1950
1951–1975
1976–2000
≥ 2000

*Figure S7*: Ancestral range probability based on the DEC model and 10,000 biogeographical stochastic mappings, plotted against the MCC species tree of Brassolini. A) The most probable state accounting for missing species in the posterior phylogenies is plotted on every node. B) The probabilities of ancestral ranges accounting for missing species in the posterior phylogenies are as pie charts on every node. C) The most probable state using the sampled species trees is plotted on every node. D) The probabilities of ancestral ranges using the sampled species trees are as pie charts on every node. Bioregions were coded as follows: M: Mesoamerica and Chocó, S: Amazonia, C: South American dry diagonal, F: Brazilian Atlantic Forest.

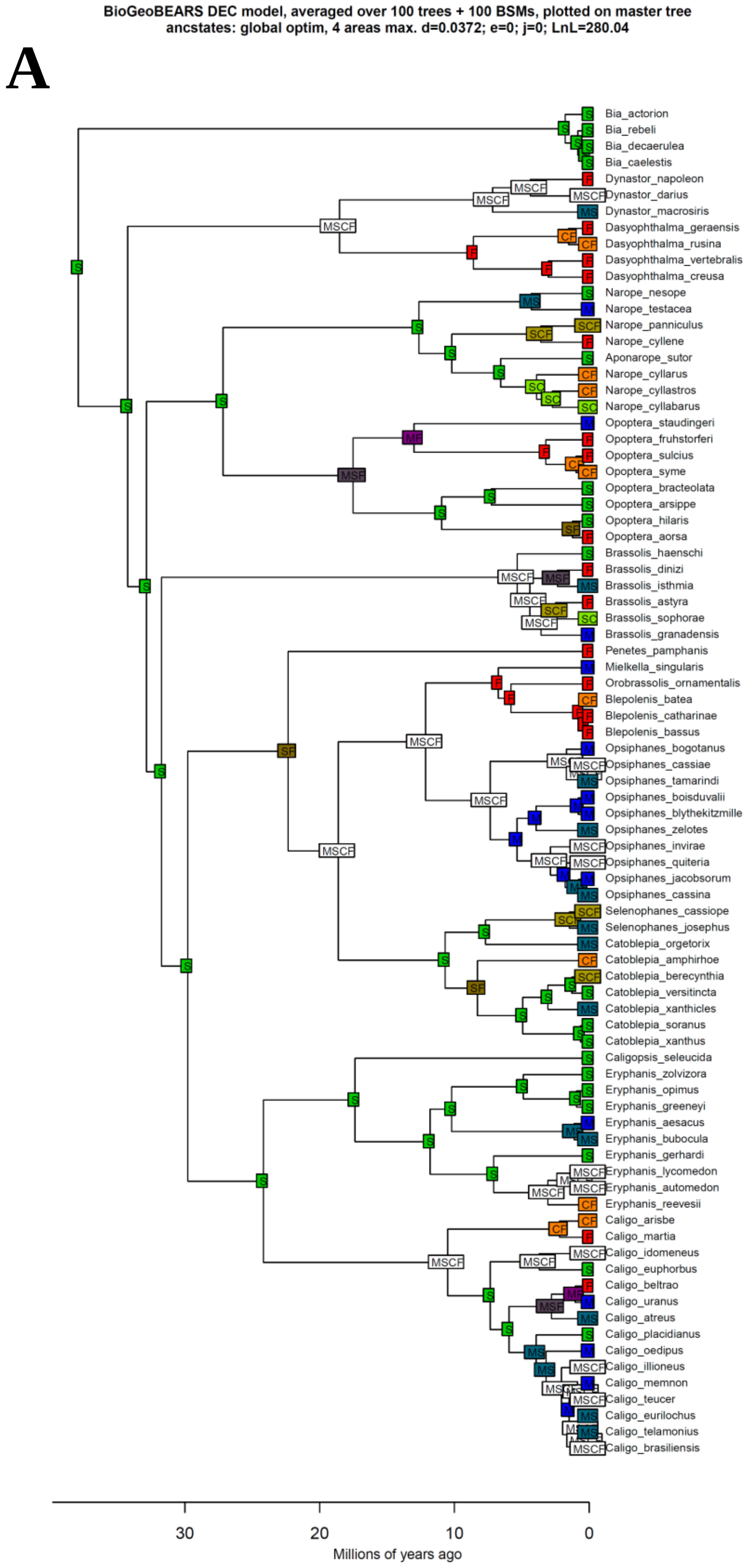
BioGeoBEARS DEC model, averaged over 100 trees + 100 BSMs, plotted on master tree
ancstates: global optim, 4 areas max. d=0.0372; e=0; j=0; LnL=280.04
A
Bia_actorion
Bia_rebeli
Bia_decaerulea
Bia_caelestis
Dynastor_napoleon
Dynastor_darius
Dynastor_macrosiris
Dasyophthalma_geraensis
Dasyophthalma_rusina
Dasyophthalma_vertebralis
Dasyophthalma_creusa
Narope_nesope
Narope_testacea
Narope_panniculus
Narope_cyllene
Aponarope_sutor
Narope_cyllarus
Narope_cyllastros
Narope_cyllabarus
Opoptera_staudingeri
Opoptera_fruhstorferi
Opoptera_sulcius
Opoptera_syme
Opoptera_bracteolata
Opoptera_arsippe
Opoptera_hilaris
Opoptera_aorsa
Brassolis_haenschi
Brassolis_dinizi
Brassolis_isthmia
Brassolis_astyra
Brassolis_sophorae
Brassolis_granadensis
Penetes_pamphanis
Mielkella_singularis
Orobrassolis_ornamentalis
Blepolenis_batea
Blepolenis_catharinae
Blepolenis_bassus
Opsiphanes_bogotanus
Opsiphanes_cassiae
Opsiphanes_tamarindi
Opsiphanes_boisduvalii
Opsiphanes_blythekitzmille
Opsiphanes_zelotes
Opsiphanes_invirae
Opsiphanes_quiteria
Opsiphanes_jacobsorum
Opsiphanes_cassina
Selenophanes_cassiope
Selenophanes_josephus
Catoblepia_orgetorix
Catoblepia_amphirhoe
Catoblepia_berecynthia
Catoblepia_versitincta
Catoblepia_xanthicles
Catoblepia_soranus
Catoblepia_xanthus
Caligopsis_seleucida
Eryphanis_zolvizora
Eryphanis_opimus
Eryphanis_greeneyi
Eryphanis_aesacus
Eryphanis_bubocula
Eryphanis_gerhardi
Eryphanis_lycomedon
Eryphanis_automedon
Eryphanis_reevesii
Caligo_arisbe
Caligo_martia
Caligo_idomeneus
Caligo_euphorbus
Caligo_beltrao
Caligo_uranus
Caligo_atreus
Caligo_placidianus
Caligo_oedipus
Caligo_illioneus
Caligo_memnon
Caligo_teucer
Caligo_eurilochus
Caligo_telamonius
Caligo_brasiliensis
30
20
10
0
Millions of years ago

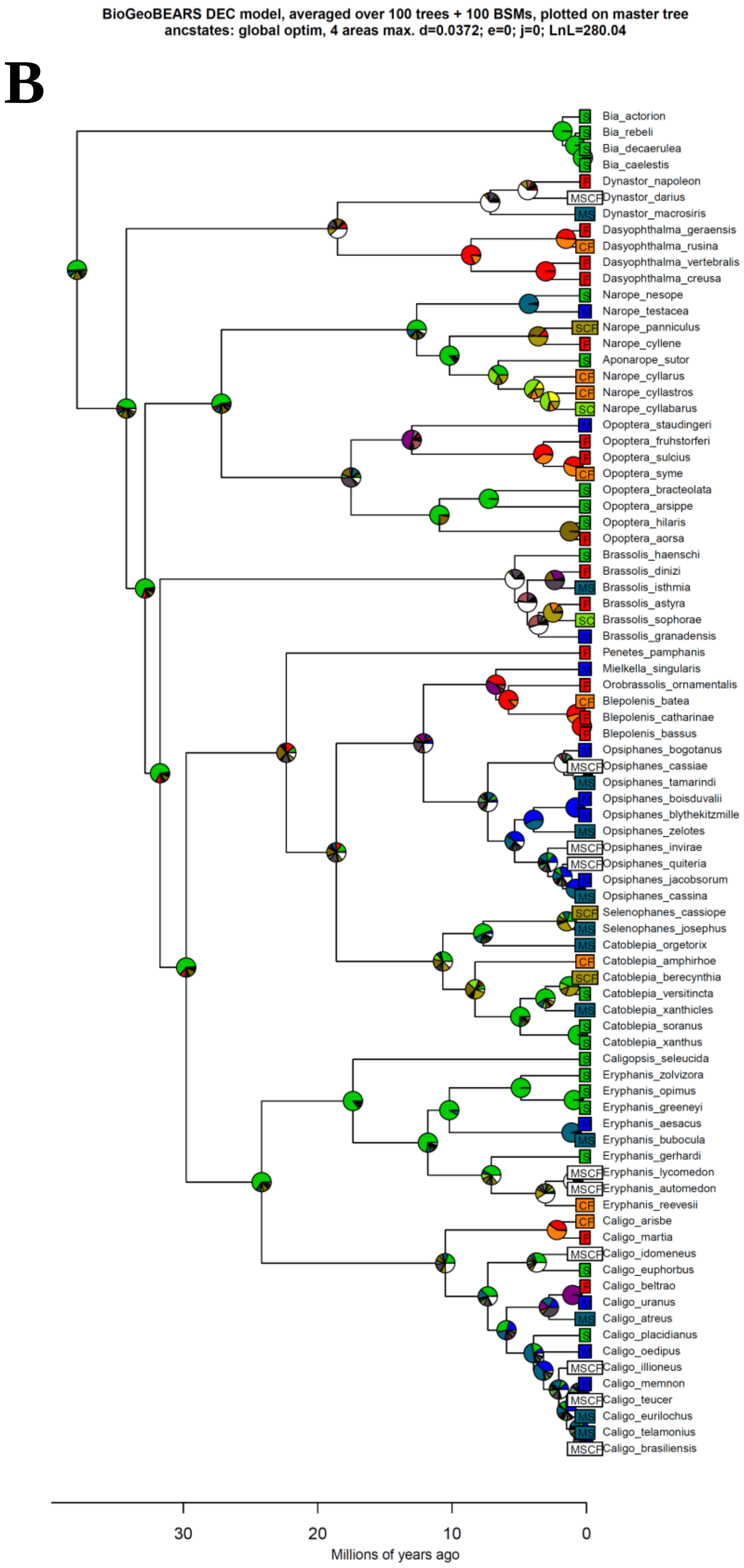
BioGeoBEARS DEC model, averaged over 100 trees + 100 BSMs, plotted on master tree
ancstates: global optim, 4 areas max. d=0.0372; e=0; j=0; LnL=280.04
B
Bia_actorion
Bia_rebeli
Bia_decaerulea
Bia_caelestis
Dynastor_napoleon
Dynastor_darius
Dynastor_macrosiris
Dasyophthalma_geraensis
Dasyophthalma_rusina
Dasyophthalma_vertebralis
Dasyophthalma_creusa
Narope_nesope
Narope_testacea
Narope_panniculus
Narope_cyllene
Aponarope_sutor
Narope_cyllarus
Narope_cyllastros
Narope_cyllabarus
Opoptera_staudingeri
Opoptera_fruhstorferi
Opoptera_sulcius
Opoptera_syme
Opoptera_bracteolata
Opoptera_arsippe
Opoptera_hilaris
Opoptera_aorsa
Brassolis_haenschi
Brassolis_dinizi
Brassolis_isthmia
Brassolis_astyra
Brassolis_sophorae
Brassolis_granadensis
Penetes_pamphanis
Mielkella_singularis
Orobrassolis_ornamentalis
Blepolenis_batea
Blepolenis_catharinae
Blepolenis_bassus
Opsiphanes_bogotanus
Opsiphanes_cassiae
Opsiphanes_tamarindi
Opsiphanes_boisduvalii
Opsiphanes_blythekitzmille
Opsiphanes_zelotes
Opsiphanes_invirae
Opsiphanes_quiteria
Opsiphanes_jacobsorum
Opsiphanes_cassina
Selenophanes_cassiope
Selenophanes_josephus
Catoblepia_orgetorix
Catoblepia_amphirhoe
Catoblepia_berecynthia
Catoblepia_versitincta
Catoblepia_xanthicles
Catoblepia_soranus
Catoblepia_xanthus
Caligopsis_seleucida
Eryphanis_zolvizora
Eryphanis_opimus
Eryphanis_greeneyi
Eryphanis_aesacus
Eryphanis_bubocula
Eryphanis_gerhardi
Eryphanis_lycomedon
Eryphanis_automedon
Eryphanis_reevesii
Caligo_arisbe
Caligo_martia
Caligo_idomeneus
Caligo_euphorbus
Caligo_beltrao
Caligo_uranus
Caligo_atreus
Caligo_placidianus
Caligo_oedipus
Caligo_illioneus
Caligo_memnon
Caligo_teucer
Caligo_eurilochus
Caligo_telamonius
Caligo_brasiliensis
30
20
10
0
Millions of years ago

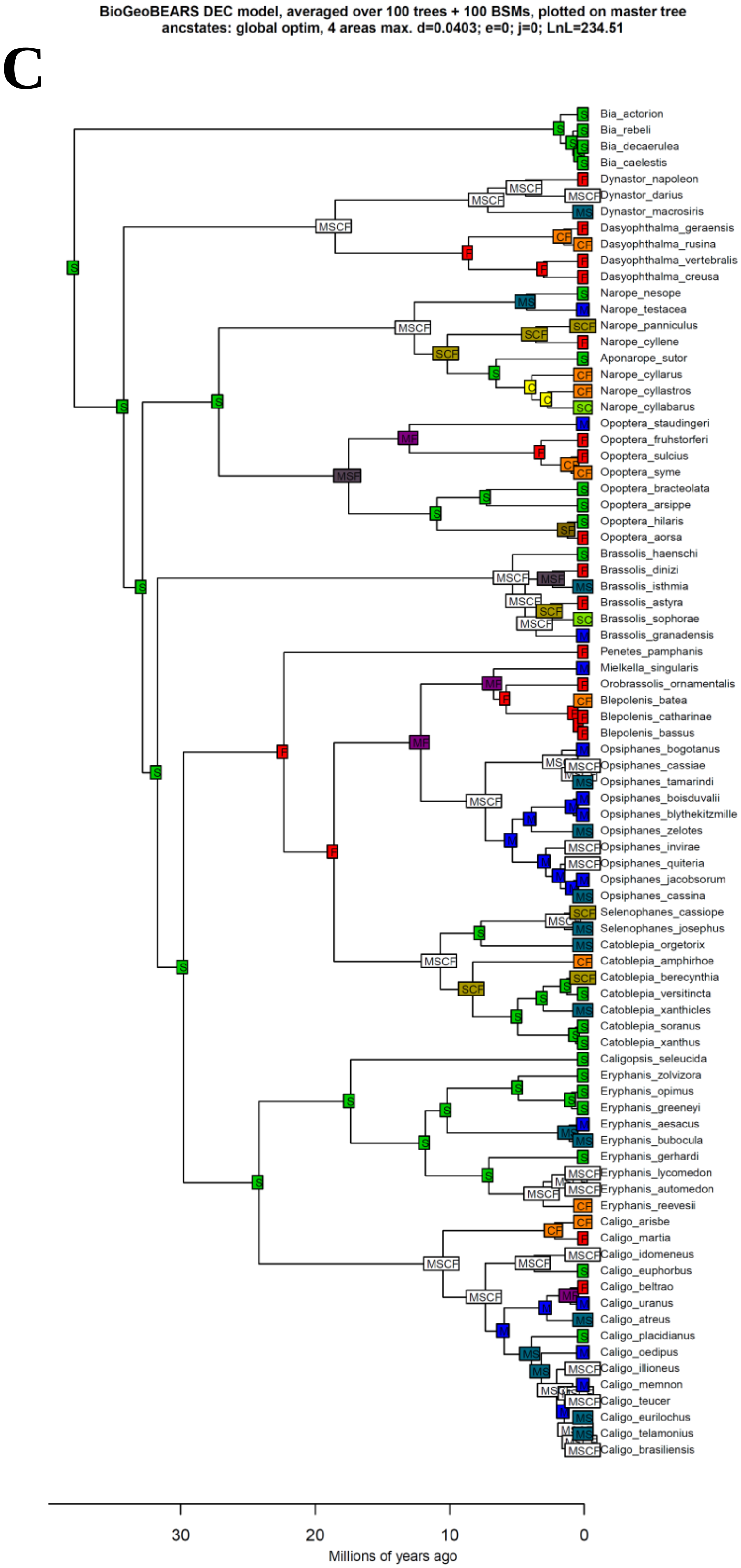
BioGeoBEARS DEC model, averaged over 100 trees + 100 BSMs, plotted on master tree
ancstates: global optim, 4 areas max. d=0.0403; e=0; j=0; LnL=234.51
C
Bia_actorion
Bia_rebeli
Bia_decaerulea
Bia_caelestis
Dynastor_napoleon
Dynastor_darius
Dynastor_macrosiris
Dasyophthalma_geraensis
Dasyophthalma_rusina
Dasyophthalma_vertebralis
Dasyophthalma_creusa
Narope_nesope
Narope_testacea
Narope_panniculus
Narope_cyllene
Aponarope_sutor
Narope_cyllarus
Narope_cyllastros
Narope_cyllabarus
Opoptera_staudingeri
Opoptera_fruhstorferi
Opoptera_sulcius
Opoptera_syme
Opoptera_bracteolata
Opoptera_arsippe
Opoptera_hilaris
Opoptera_aorsa
Brassolis_haenschi
Brassolis_dinizi
Brassolis_isthmia
Brassolis_astyra
Brassolis_sophorae
Brassolis_granadensis
Penetes_pamphanis
Mielkella_singularis
Orobrassolis_ornamentalis
Blepolenis_batea
Blepolenis_catharinae
Blepolenis_bassus
Opsiphanes_bogotanus
Opsiphanes_cassiae
Opsiphanes_tamarindi
Opsiphanes_boisduvalii
Opsiphanes_blythekitzmille
Opsiphanes_zelotes
Opsiphanes_invirae
Opsiphanes_quiteria
Opsiphanes_jacobsorum
Opsiphanes_cassina
Selenophanes_cassiope
Selenophanes_josephus
Catoblepia_orgetorix
Catoblepia_amphirhoe
Catoblepia_berecynthia
Catoblepia_versitincta
Catoblepia_xanthicles
Catoblepia_soranus
Catoblepia_xanthus
Caligopsis_seleucida
Eryphanis_zolvizora
Eryphanis_opimus
Eryphanis_greeneyi
Eryphanis_aesacus
Eryphanis_bubocula
Eryphanis_gerhardi
Eryphanis_lycomedon
Eryphanis_automedon
Eryphanis_reevesii
Caligo_arisbe
Caligo_martia
Caligo_idomeneus
Caligo_euphorbus
Caligo_beltrao
Caligo_uranus
Caligo_atreus
Caligo_placidianus
Caligo_oedipus
Caligo_illioneus
Caligo_memnon
Caligo_teucer
Caligo_eurilochus
Caligo_telamonius
Caligo_brasiliensis
30
20
10
0
Millions of years ago

D

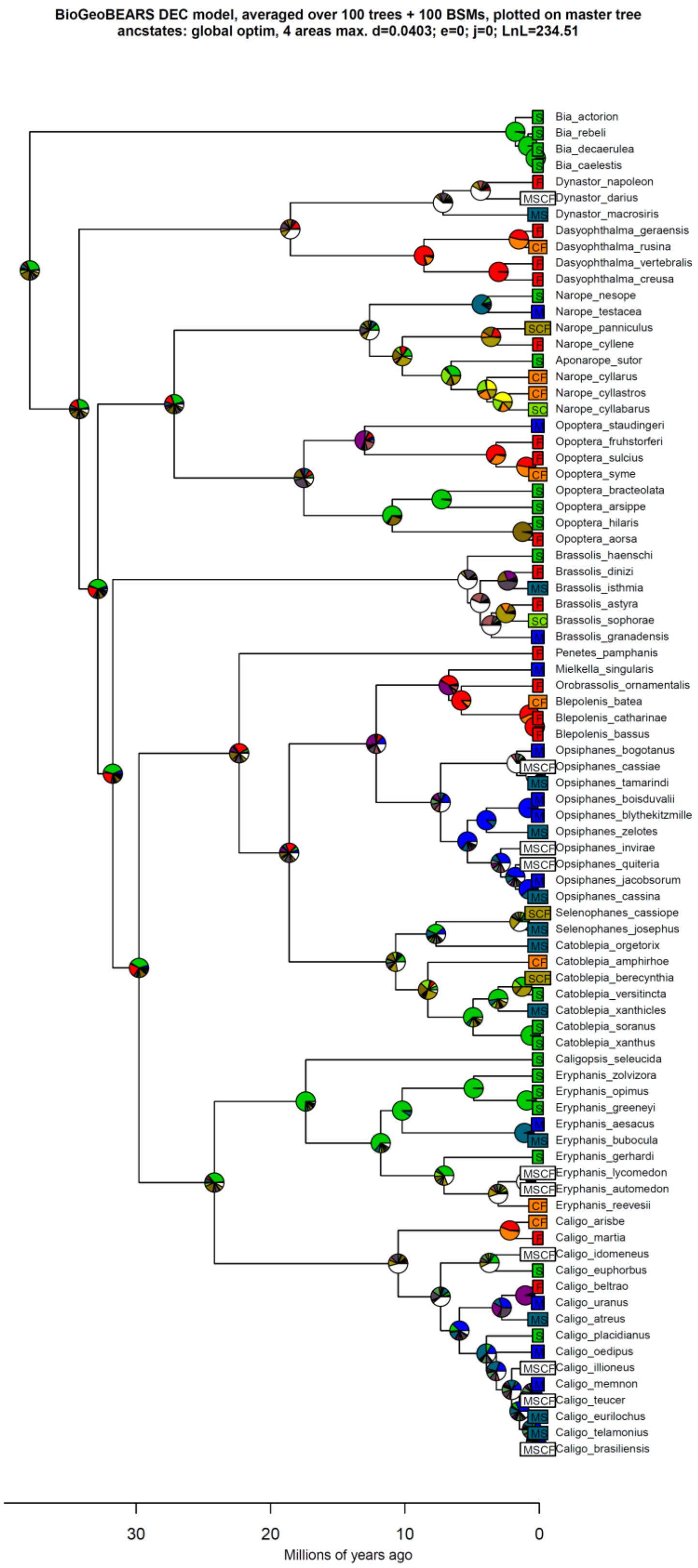

BioGeoBEARS DEC model, averaged over 100 trees + 100 BSMs, plotted on master tree
ancstates: global optim, 4 areas max. d=0.0403; e=0; j=0; LnL=234.51
Bia_actorion
Bia_rebeli
Bia_decaerulea
Bia_caelestis
Dynastor_napoleon
Dynastor_darius
Dynastor_macrosiris
Dasyophthalma_geraensis
Dasyophthalma_rusina
Dasyophthalma_vertebralis
Dasyophthalma_creusa
Narope_nesope
Narope_testacea
Narope_panniculus
Narope_cyllene
Aponarope_sutor
Narope_cyllarus
Narope_cyllastros
Narope_cyllabarus
Opoptera_staudingeri
Opoptera_fruhstorferi
Opoptera_sulcius
Opoptera_syme
Opoptera_bracteolata
Opoptera_arsippe
Opoptera_hilaris
Opoptera_aorsa
Brassolis_haenschi
Brassolis_dinizi
Brassolis_isthmia
Brassolis_astyra
Brassolis_sophorae
Brassolis_granadensis
Penetes_pamphanis
Mielkella_singularis
Orobrassolis_ornamentalis
Blepolenis_batea
Blepolenis_catharinae
Blepolenis_bassus
Opsiphanes_bogotanus
Opsiphanes_cassiae
Opsiphanes_tamarindi
Opsiphanes_boisduvalii
Opsiphanes_blythekitzmille
Opsiphanes_zelotes
Opsiphanes_invirae
Opsiphanes_quiteria
Opsiphanes_jacobsorum
Opsiphanes_cassina
Selenophanes_cassiope
Selenophanes_josephus
Catoblepia_orgetorix
Catoblepia_amphirhoe
Catoblepia_berecynthia
Catoblepia_versitincta
Catoblepia_xanthicles
Catoblepia_soranus
Catoblepia_xanthus
Caligopsis_seleucida
Eryphanis_zolvizora
Eryphanis_opimus
Eryphanis_greeneyi
Eryphanis_aesacus
Eryphanis_bubocula
Eryphanis_gerhardi
Eryphanis_lycomedon
Eryphanis_automedon
Eryphanis_reevesii
Caligo_arisbe
Caligo_martia
Caligo_idomeneus
Caligo_euphorbus
Caligo_beltrao
Caligo_uranus
Caligo_atreus
Caligo_placidianus
Caligo_oedipus
Caligo_illioneus
Caligo_memnon
Caligo_teucer
Caligo_eurilochus
Caligo_telamonius
Caligo_brasiliensis
30
20
10
0
Millions of years ago

*Figure S8*: Dispersal rates through time calculated with 10,000 biogeographical stochastic mappings in BioGeoBEARS. Bioregions were coded as follows: M: Mesoamerica and Chocó, S: Amazonia, C: South American dry diagonal, F: Brazilian Atlantic Forest. The x axis in every chart is at million years scale. The y axis represents the estimated dispersal rates (events per lineage per million years) using the formula in Antonelli et al. (2018). “rate.StoM”, for example, is dispersal from source area “S” to target area “M”. Solid lines are the median values, dark green ribbons represent the lower and upper quartiles (0.25 and 0.75 quantiles), light green ribbons the 0.1 and 0.9 quantiles, and dashed lines the 0.05 and 0.95 quantiles. A) Estimates accounting for missing species in the posterior phylogenies. B) Estimates based on the sampled species trees.

**A**

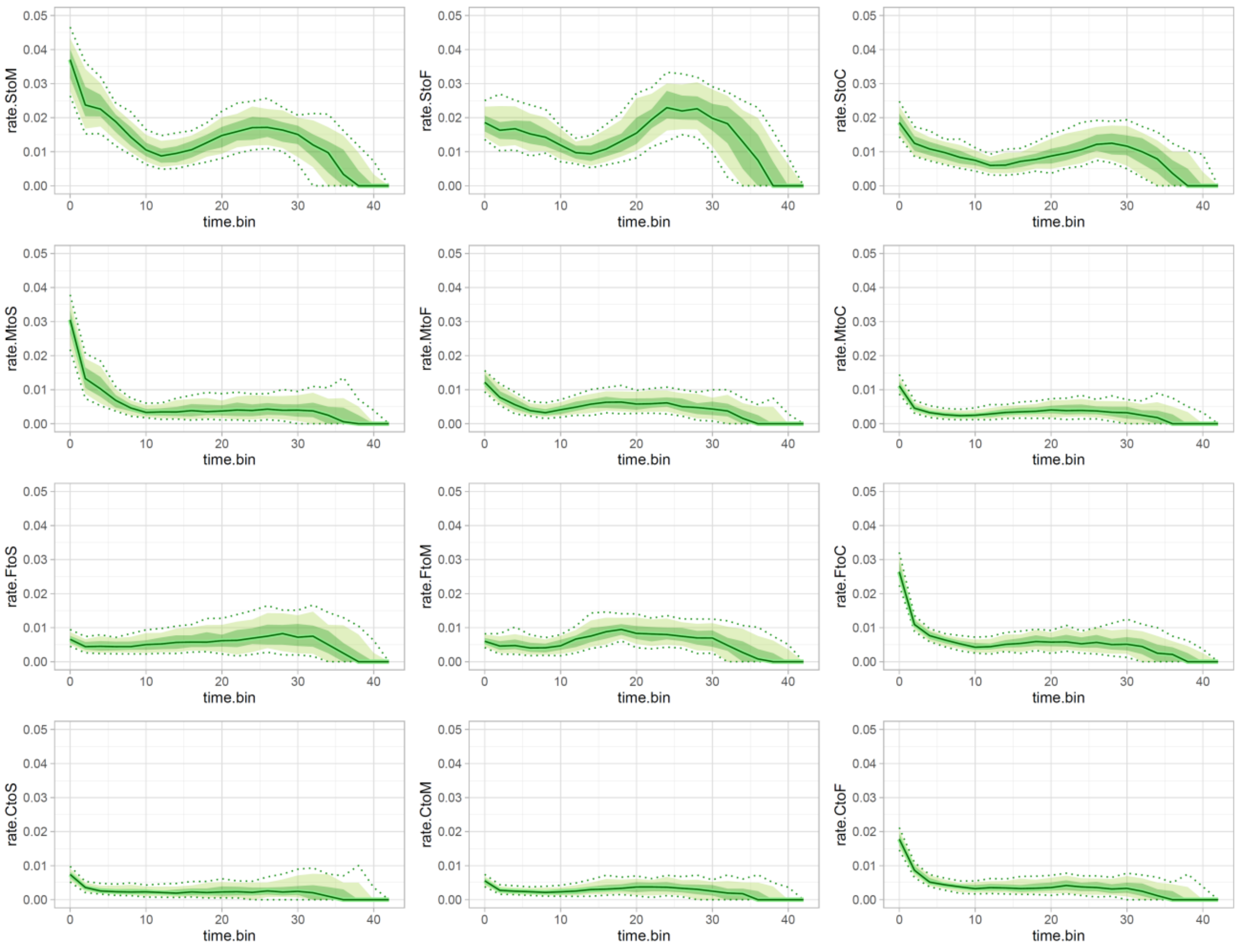
rate.StoM
rate.StoF
rate.StoC
rate.MtoS
rate.MtoF
rate.MtoC
rate.FtoS
rate.FtoM
rate.FtoC
rate.CtoS
rate.CtoM
rate.CtoF
time.bin
0.00
0.01
0.02
0.03
0.04
0.05
0
10
20
30
40

# B

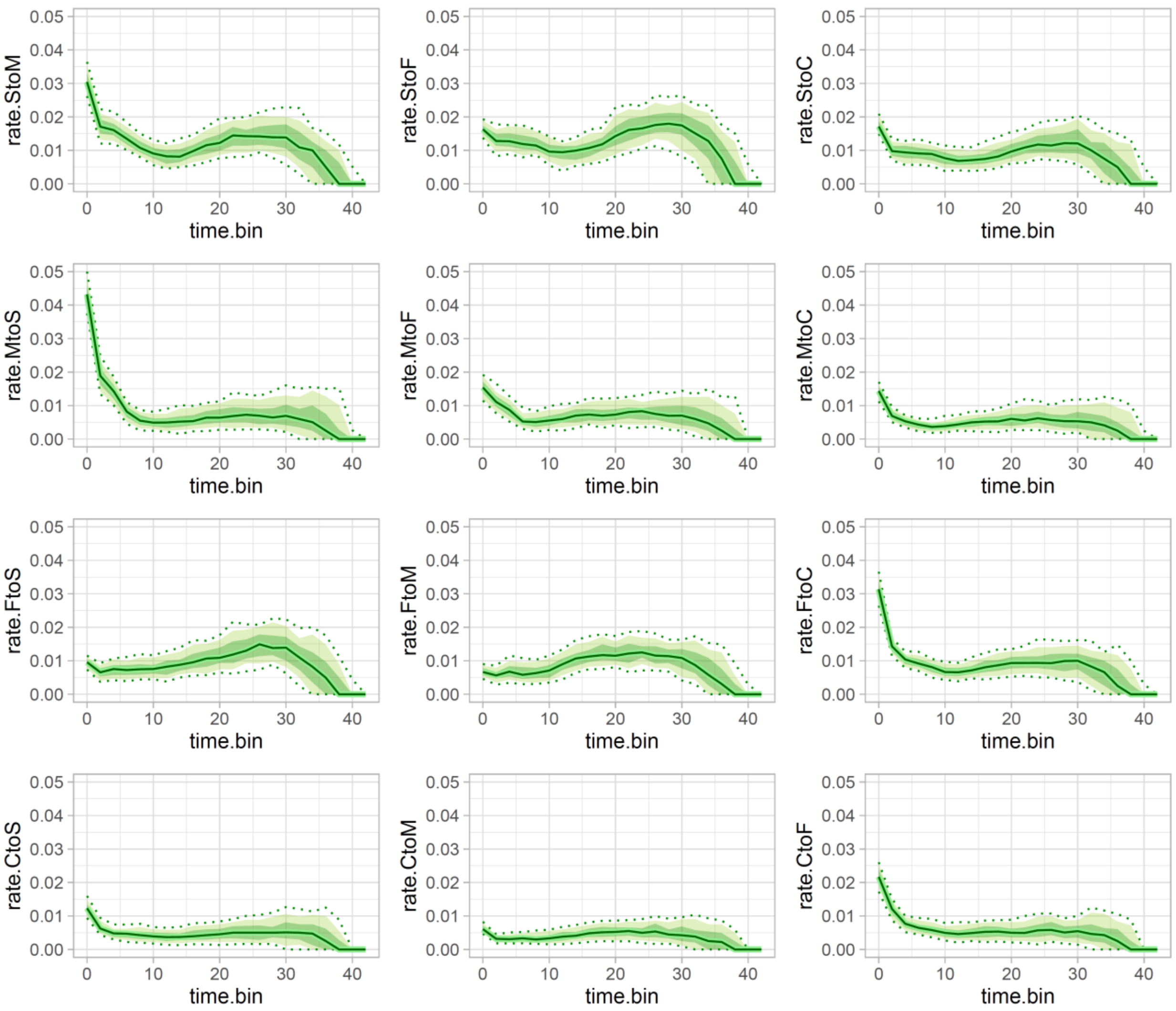
rate.StoM
rate.StoF
rate.StoC
rate.MtoS
rate.MtoF
rate.MtoC
rate.FtoS
rate.FtoM
rate.FtoC
rate.CtoS
rate.CtoM
rate.CtoF
time.bin
0.00 0.01 0.02 0.03 0.04 0.05
0 10 20 30 40

*Figure S9*: Within-area cladogenesis through time calculated with 10,000 biogeographical stochastic mappings in BioGeoBEARS. Bioregions were coded as follows: M: Mesoamerica and Chocó, S: Amazonia, C: South American dry diagonal, F: Brazilian Atlantic Forest. The x axis in every chart is at million years scale. The y axis represents the estimated relative number of cladogenesis events per million years using a formula modified from Xing and Ree (2017). “rate.StoS”, for example, is relative *in situ* cladogenesis in area “S”. Solid lines are the median values, dark green ribbons represent the lower and upper quartiles (0.25 and 0.75 quantiles), light green ribbons the 0.1 and 0.9 quantiles, and dashed lines the 0.05 and 0.95 quantiles. A) Estimates accounting for missing species in the posterior phylogenies. B) Estimates based on the sampled species trees.

**A**

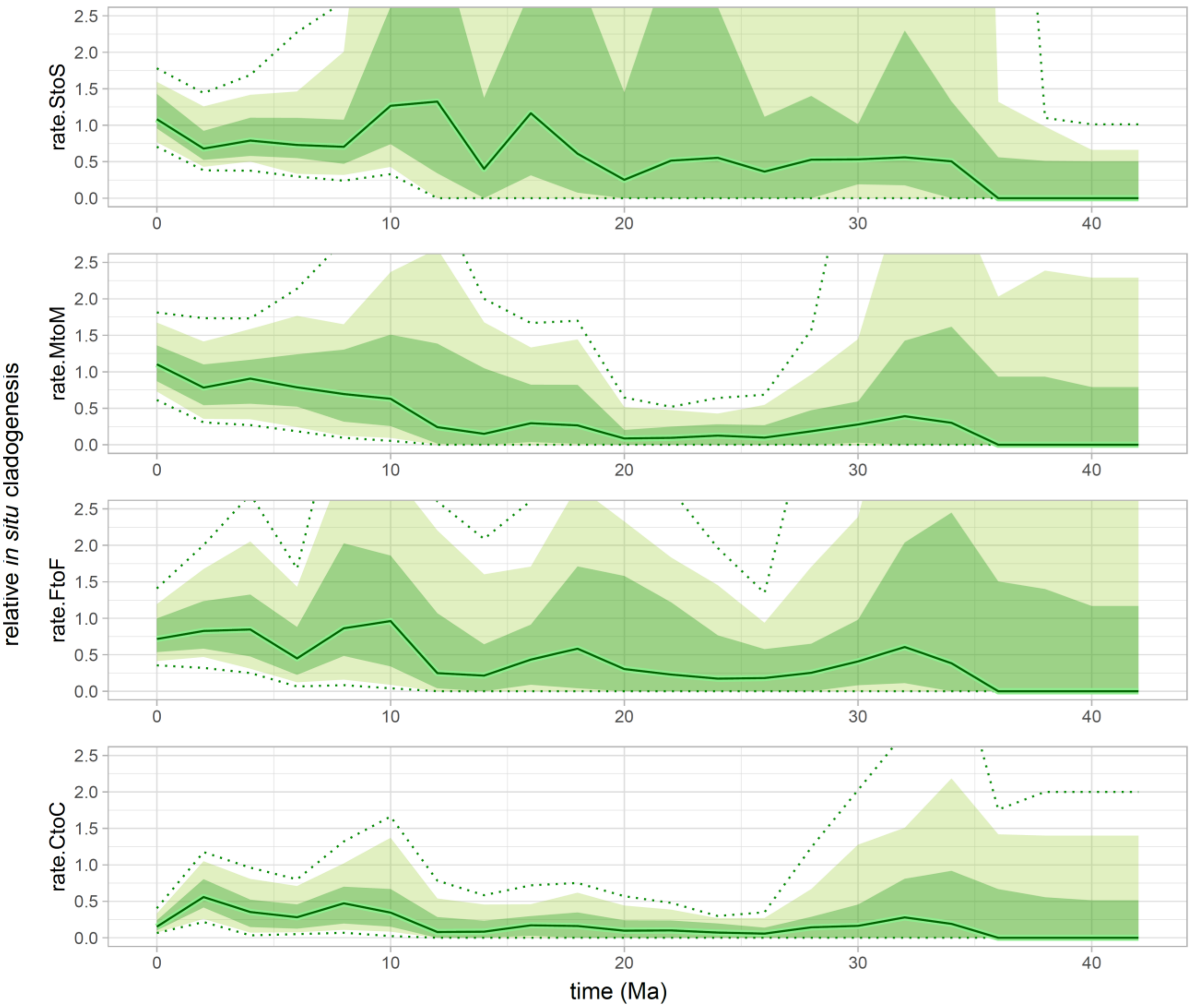

relative *in situ* cladogenesis
rate.StoS
rate.MtoM
rate.FtoF
rate.CtoC
2.5
2.0
1.5
1.0
0.5
0.0
0
10
20
30
40
time (Ma)

**B**

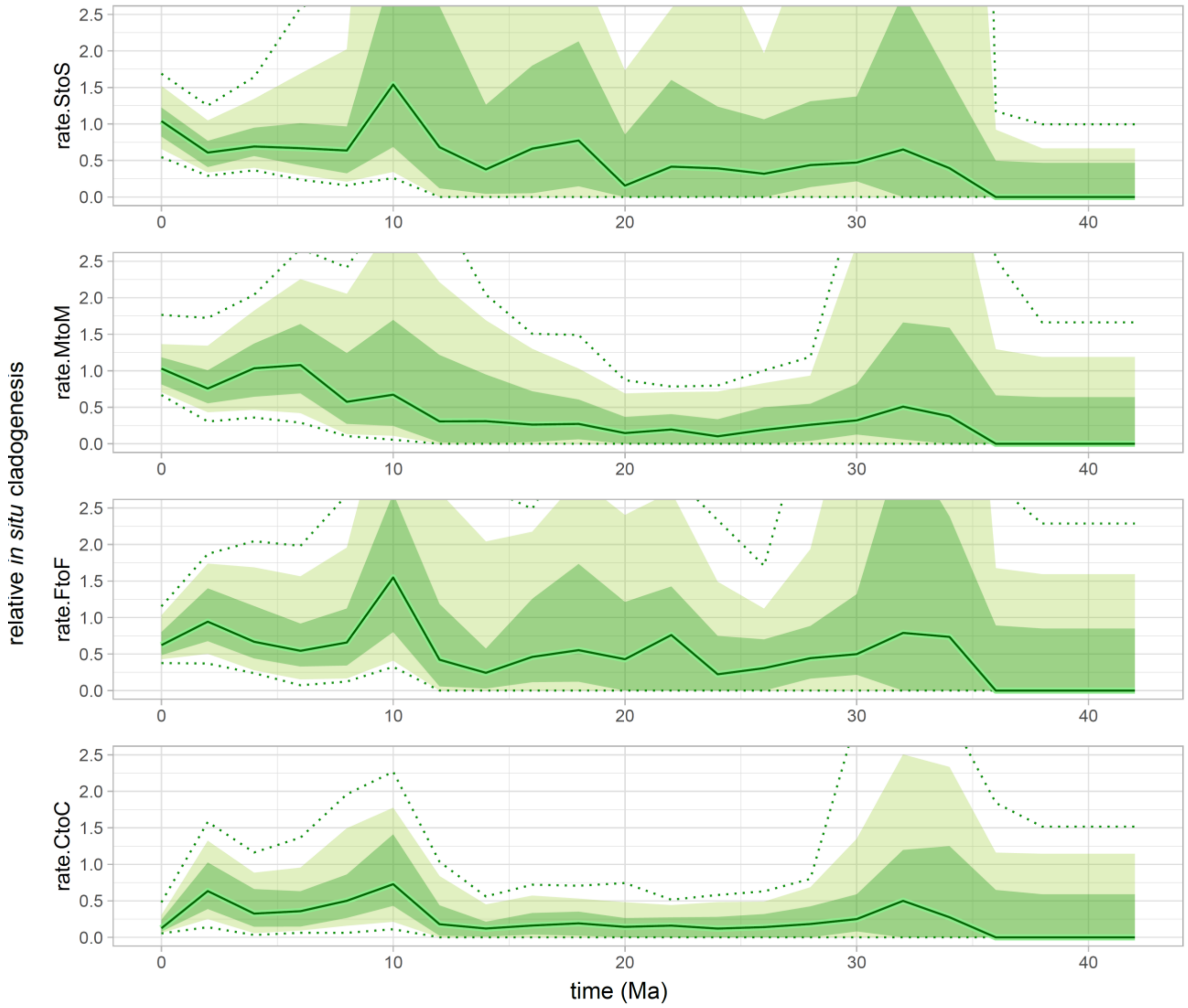

relative *in situ* cladogenesis
rate.StoS
rate.MtoM
rate.FtoF
rate.CtoC
0.0
0.5
1.0
1.5
2.0
2.5
0
10
20
30
40
time (Ma)

*Table S1* (As a separate file): Voucher locality information and associated genetic data deposited in GenBank or BOLD (ASARD codes).

*Table S2*: The best-fit partition scheme for the molecular dataset estimated by the program PartitionFinder v.2.1.1.

| **Subset** | **N° of sites** | **Partition names** |
|---|---|---|
| 1 | 284 | CAD_pos3 |
| 2 | 420 | CAD_pos1, wingless_pos1 |
| 3 | 1622 | EF1α_pos2, CAD_pos2, RpS5_pos2, COI_pos2, GAPDH_pos2 |
| 4 | 492 | COI_pos3 |
| 5 | 492 | COI_pos1 |
| 6 | 851 | GAPDH_pos3, RpS5_pos3, EF1α_pos3 |
| 7 | 986 | wingless_pos2, RpS5_pos1, EF1α_pos1, GAPDH_pos1 |
| 8 | 138 | wingless_pos3 |

*Table S3*: The best-fit partition scheme for the morphological dataset estimated by homoplasy scores (*f*) in the program TNT v.1.5.

| **Subset** | ***f*** | **N° characters** | **Morphological characters** |
|---|---|---|---|
| 1 | 0.00 | 103 | 3 5 16 30 32 33 34 36 46 49 54 58 67 69 79 81 83 91 94 97 101 103 104 105 109 114 116 117 118 128 132 134 136 137 138 140 142 144 149 150 152 154 155 156 161 162 163 164 165 166 170 171 174 176 178 179 180 181 182 184 186 188 189 190 191 194 196 198 199 203 205 206 208 209 210 212 214 215 216 217 218 220 223 224 225 227 230 232 233 235 236 237 238 239 240 243 244 248 250 252 253 254 255 |
| 2 | 0.25 | 40 | 1 7 14 27 42 44 62 63 66 80 87 99 107 108 110 111 112 121 123 124 125 135 145 151 153 157 160 168 172 177 200 202 204 207 213 221 229 231 247 249 |
| 3 | 0.40 | 26 | 9 12 21 37 38 40 48 57 71 73 74 86 93 96 119 120 127 129 131 133 159 222 228 245 246 251 |
| 4 | 0.50 | 12 | 10 15 17 18 28 43 47 56 60 77 106 115 |
| 5 | 0.57 | 5 | 51 53 59 98 242 |

| | | | |
|---|---|---|---|
| 6 | 0.63 | 9 | 19 22 24 31 35 41 50 72 241 |
| 7 | 0.67 | 8 | 11 20 23 26 29 55 65 130 |
| 8 | 0.70 | 2 | 39 78 |
| 9 | 0.73 | 2 | 52 70 |
| 10 | 0.75 | 1 | 45 |
| 11 | non-informative | 47 | 2 4 6 8 13 25 61 64 68 75 76 82 84 85 88 89 90 92 95 100 102 113 122 126 139 141 143 146 147 148 158 167 169 173 175 183 185 187 192 193 195 197 201 211 219 226 234 |

*Table S4*: Bayes factor comparison between the strict and relaxed clock models. SSML: Stepping-stone marginal likelihood; BF: Bayes factor calculated as twice its natural logarithm (2 $\log_e BF$), and to account for the number of parameters (NP), we summed to this value the following: $(NP_{relaxed} - NP_{strict}) \times \log_e 0.01$. The relaxed clock model was strongly preferred over the strict clock for all loci (BF > 10).

| **Molecular clock** | **Dataset** | **N° parameters** | **SSML** | **BF** |
|---|---|---|---|---|
| Relaxed | COI | 15 | −23762.54 | 25.99 |
| *Strict* | *COI* | *13* | *−23780.14* | — |
| Relaxed | CAD | 15 | −6797.52 | 48.97 |
| *Strict* | *CAD* | *13* | *−6826.61* | — |
| Relaxed | EF1α | 15 | −12996.34 | 398.77 |
| *Strict* | *EF1α* | *13* | *−13200.33* | — |
| Relaxed | GAPDH | 15 | −7014.36 | 11.05 |
| *Strict* | *GAPDH* | *13* | *−7024.49* | — |
| Relaxed | RpS5 | 15 | −7861.07 | 116.51 |
| *Strict* | *RpS5* | *13* | *−7923.93* | — |
| Relaxed | *wingless* | 15 | −5417.38 | 142.41 |
| *Strict* | *wingless* | *13* | *−5493.19* | — |

*Table S5*: Bayes factor comparisons among six models involving two tree models (Yule and birth-death) and molecular clock partitions (one, two, mitochondrial and nuclear, and six, for each gene partition). Marginal L (likelihood) estimates based on 25 path-sampling steps under thermodynamic integration. BF: Bayes factor with respect to the highest-likelihood model (Yule + 2 clocks) calculated as twice its natural logarithm (2 $\log_e$ *BF*). The Yule + 2 molecular clocks model received decisive support (BF > 10).

| **Tree model** | **Molecular clocks** | **Marginal L estimate** | **BF** |
|---|---|---|---|
| *Yule* | 2 | −64584.43 | — |
| *Yule* | 6 | −64607.00 | −45.14 |
| *Birth-death* | 6 | −64614.35 | −59.82 |
| *Birth-death* | 2 | −64691.74 | −214.62 |
| *Birth-death* | 1 | −64875.58 | −582.30 |
| *Yule* | 1 | −64903.45 | −638.04 |

*Table S6*: Tree topology test comparing the branching order of early divergent Brassolini lineages. The total-evidence concatenation-based consensus tree (*Conc*) has the following constraint topology (*Bia*: ((*Dasyophthalma*:*Dynastor*): ((*Narope*:*Opoptera*): ((*Caligo*-group): ((*Brassolis*: *Opsiphanes*-group)))))), which is highly-similar to the unconstrained molecular tree (*Unconstr*). The multispecies coalescent tree (*MSC*) has the following constraint topology (*Bia*: ((*Dasyophthalma*:*Dynastor*): ((*Narope*:*Opoptera*): (Brassolis: (*Caligo*-group: *Opsiphanes*-group))))). The morphology-based systematics of Brassolini (*Naropina*) has the following constraint topology (*Bia*: (*Narope*: remaining Brassolina genera)). DeltaL: Likelihood difference from the maximum likelihood tree topology; bp-RELL: bootstrap proportion using the RELL method; p-KH: p-value of one sided Kishino-Hasegawa test; p-SH: p-value of Shimodaira-Hasegawa test; p-WKH: p-value of weighted KH test; p-WSH: p-value of weighted SH test; c-ELW: expected likelihood weight; p-AU: p-value of approximately unbiased test. All tests performed 10,000 resamplings using the RELL method. The plus signs denote the 95% confidence sets. The minus signs denote significant exclusion. The tested branching orders are undecidable given our molecular dataset, except for *Naropina*, which is significantly rejected.

| **Tree** | **LogL** | **DeltaL** | **bp-RELL** | **p-KH** | **p-SH** | **p-WKH** | **p-WSH** | **c-ELW** | **p-AU** |
|---|---|---|---|---|---|---|---|---|---|
| *Unconstr* | −58956.83556 | 0 | 0.416 + | 0.775 + | 1 + | 0.775 + | 0.988 + | 0.435 + | 0.87 + |
| *Conc* | −58956.86496 | 0.029398 | 0.428 + | 0.225 + | 0.821 + | 0.225 + | 0.556 + | 0.422 + | 0.296 + |
| *MSC* | −58962.88425 | 6.0487 | 0.156 + | 0.155 + | 0.411 + | 0.155 + | 0.255 + | 0.143 + | 0.119 + |
| *Naropina* | −59012.47309 | 55.638 | 0.0001 − | 0.0001 − | 0.0004 − | 0.0001 − | 0.0001 − | 7.97*e*−05 − | 1.59*e*−05 − |

*Table S7*: Sampling fractions used for taking into account missing species in the species tree for calculation of dispersal and speciation rates. The revised genera are monophyletic, and, given our comprehensive taxonomic sampling, we assumed that missing taxa are within crown nodes.

| **Genus** | **Described species** | **Sampled species** | **Fraction sampled** |
|---|---:|---:|---:|
| *Bia* | 6 | 4 | 0.6667 |
| *Blepolenis* | 3 | 3 | 1.0000 |
| *Brassolis* | 6 | 6 | 1.0000 |
| *Caligo* | 22 | 15 | 0.6818 |
| *Caligopsis* | 1 | 1 | 1.0000 |
| *Catoblepia* | 7 | 6 | 0.8571 |
| *Dasyophthalma* | 4 | 4 | 1.0000 |
| *Dynastor* | 3 | 3 | 1.0000 |
| *Eryphanis* | 9 | 9 | 1.0000 |
| *Mielkella* | 1 | 1 | 1.0000 |
| *Narope* | 18 | 8 | 0.4444 |
| *Opoptera* | 8 | 8 | 1.0000 |
| *Opsiphanes* | 13 | 10 | 0.7692 |
| *Orobrassolis* | 2 | 1 | 0.5000 |
| *Penetes* | 1 | 1 | 1.0000 |
| *Selenophanes* | 3 | 2 | 0.6667 |
